# The pluripotency factor Tex10 finetunes Wnt signaling for PGC and male germline development

**DOI:** 10.1101/2023.02.23.529824

**Authors:** Dan Li, Jihong Yang, Fanglin Ma, Vikas Malik, Ruge Zang, Xianle Shi, Xin Huang, Hongwei Zhou, Jianlong Wang

## Abstract

Testis-specific transcript 10 (Tex10) is a critical factor for pluripotent stem cell maintenance and preimplantation development. Here, we dissect its late developmental roles in primordial germ cell (PGC) specification and spermatogenesis using cellular and animal models. We discover that Tex10 binds the Wnt negative regulator genes, marked by H3K4me3, at the PGC-like cell (PGCLC) stage in restraining Wnt signaling. Depletion and overexpression of Tex10 hyperactivate and attenuate the Wnt signaling, resulting in compromised and enhanced PGCLC specification efficiency, respectively. Using the Tex10 conditional knockout mouse models combined with single-cell RNA sequencing, we further uncover critical roles of Tex10 in spermatogenesis with Tex10 loss causing reduced sperm number and motility associated with compromised round spermatid formation. Notably, defective spermatogenesis in Tex10 knockout mice correlates with aberrant Wnt signaling upregulation. Therefore, our study establishes Tex10 as a previously unappreciated player in PGC specification and male germline development by fine-tuning Wnt signaling.

## INTRODUCTION

Infertility is caused by various health problems, such as cancer (Poorvu et al. 2019), obesity (Broughton and Moley 2017; Craig et al. 2017), and genetic mutations (Krausz and Riera-Escamilla 2018; Beke 2019). A reduction in the quality or quantity of germ cells produced by an individual could also significantly impact a person’s fertility and child health in the next generation. In humans, the appropriate specification and differentiation of primordial germ cells (PGCs), pioneering germ cells in the embryo, are critical to promote human reproductive health, which has affected around 10% of couples (Chen et al. 2017).

An *in vitro* PGC-like cell (PGCLC) model (Hayashi et al. 2011) was developed to facilitate molecular studies of PGC specification. PGCLCs are derived by first inducing differentiation of naïve embryonic stem cells (ESCs), equivalent to the inner cell mass of blastocysts (E3.5-4.5), towards competent formative epiblast-like cells (EpiLCs), transient *in vitro* cells resembling pregastrulation mouse epiblast (E5.5-6.5) (Hayashi et al. 2011). Then, EpiLCs in response to Bmp and Wnt signaling are further differentiated and specified as PGCLCs, which are equivalent to migratory PGCs *in vivo* (E8.5-10.5) and can further develop to mature functional gametes. In mice, PGC specification follows Wnt-dependent induction, which promotes the activation of several important PGC specifiers such as Prdm1 (aka Blimp1), Prdm14 and AP2γ (Tang et al. 2016). These factors further form a self-reinforcing network in promoting germ cell development and pluripotency gene reactivation while feeding back to repress other Wnt/Bmp-induced mesoderm genes (Magnusdottir et al. 2013; Tang et al. 2016). Accordingly, induction of mouse germ-cell fate can be achieved by transcription factors (TFs) such as Prdm1 and Prdm14 in vitro and such TF-induced PGCLCs can function as bona fide precursors for the spermatogenesis producing healthy offspring (Nakaki et al. 2013). Of note, both aberrant Wnt activation and inhibition impair PGCLC specification, suggesting that precise Wnt activity levels are necessary for optimal PGC specification (Hackett et al. 2018).

PGC specification in humans and mice is followed by migration, colonization, sex determination, and PGC differentiation to either sperms or oocytes (Chen et al. 2017). A notable feature of the PGC specification is the reactivation of pluripotency, a property of a cell to differentiate into three germ cell layers of the early embryo and, therefore, into all cells of the adult body. Interestingly, many pluripotency factors play an important role in the specification of PGCs, exemplified by the requirement of the key pluripotency factors Nanog (Murakami et al. 2016), Oct4 (Kehler et al. 2004), and Sox2 (Campolo et al. 2013) for PGC specification and development. In contrast, the primed pluripotency factor Otx2 plays a negative role in restricting PGC specification (Zhang et al. 2018a). Tex10 (Testis expressed 10) is a transcriptional regulator that functions as a partner of Sox2, Oct4, and Nanog in coordinating epigenetic control of super-enhancer activity in pluripotency and reprogramming (Ding et al. 2015). Loss of function studies establish the important roles of Tex10 in ESCs and preimplantation development: Tex10 knockdown ESCs cannot be maintained due to the differentiation/apoptosis, and Tex10 global knockout is early embryonic lethal at the morula to the blastocyst stage (Ding et al. 2015). Whether Tex10 plays any role in postimplantation development, particularly germ cell development, remains to be defined.

In this study, we addressed the functional contribution of Tex10 to PGC specification and male germline development. By employing the degron approach (Nabet et al. 2018), we demonstrated that the loss of Tex10 induced apoptosis and hyperactivated Wnt signaling, leading to compromised PGCLC specification efficiency. Conversely, Tex10 overexpression attenuated Wnt signaling and restricted somatic lineage programs to promote the specification of PGCLCs. Furthermore, we created a conditional knockout mouse model of Tex10 to reveal the requirement of Tex10 for male fertility, manifested by reduced sperm number and motility in male mice deficient for Tex10. Single cell transcriptome analysis further pinpointed the association of male germline defects with compromised round spermatid formation during spermatogenesis. Therefore, our study establishes a previously unappreciated role of the pluripotency factor Tex10 in PGCLC specification and spermatogenesis.

## RESULTS

### Tex10 is a candidate regulator for PGC and male germline development

Multiple lines of evidence indicate that the pluripotency factor Tex10 is specifically and highly enriched in male germline lineage. First, we curated 30 tissues/cells related to pluripotency and germ cell development from 272 distinct mouse cell types or tissues (Hutchins et al. 2017), including early embryos (such as 4-cell, 8-cell, morula, and blastocyst), ESCs, epiblast, and PGCs (male/female) and several immune cell types as reference cell types, including CD4 and CD8 T cells (Table S1). These curated samples are part of the global gene expression datasets from the collection of 921 RNA-sequencing samples (Hutchins et al. 2017) and the PGC dataset from E13.5 gonads sorted with the germ cell surface marker SSEA1 (Liu et al. 2014). Our analysis reveals that Tex10 has the highest expression in male PGCs (ranked #1) and unexpectedly, also a relatively high expression in female PGCs (ranked #5) among 30 tissues/cell lines (Fig. 1A). This germ cell enrichment of Tex10 is like that of the PGC master regulator Prdm1 (Tang et al. 2016) (ranked #1 in both male and female PGCs; Fig. S1A). Second, *Tex10* expression is gradually increased in the testes of day 1, day 3, and day 7 and in spermatocytes from adult mouse testes (Fig. 1B and Table S2). In contrast, *Tex10* did not show a consistent trend of up-regulation or down-regulation during the development of female PGC to MII oocytes (Fig. S1B and Table S3). Third, BioGPS, an online gene annotation portal, shows that Tex10 is specifically and highly enriched in testes (and ESCs) among many different cell lines and tissues (Fig. S1C). Given its prominent roles in transcriptional and epigenetic control of stem cell pluripotency, these data strongly suggest that Tex10 could be a previously unrecognized important regulator of PGCLC specification and play a preferred role in male germline development.

**Fig. 1.**
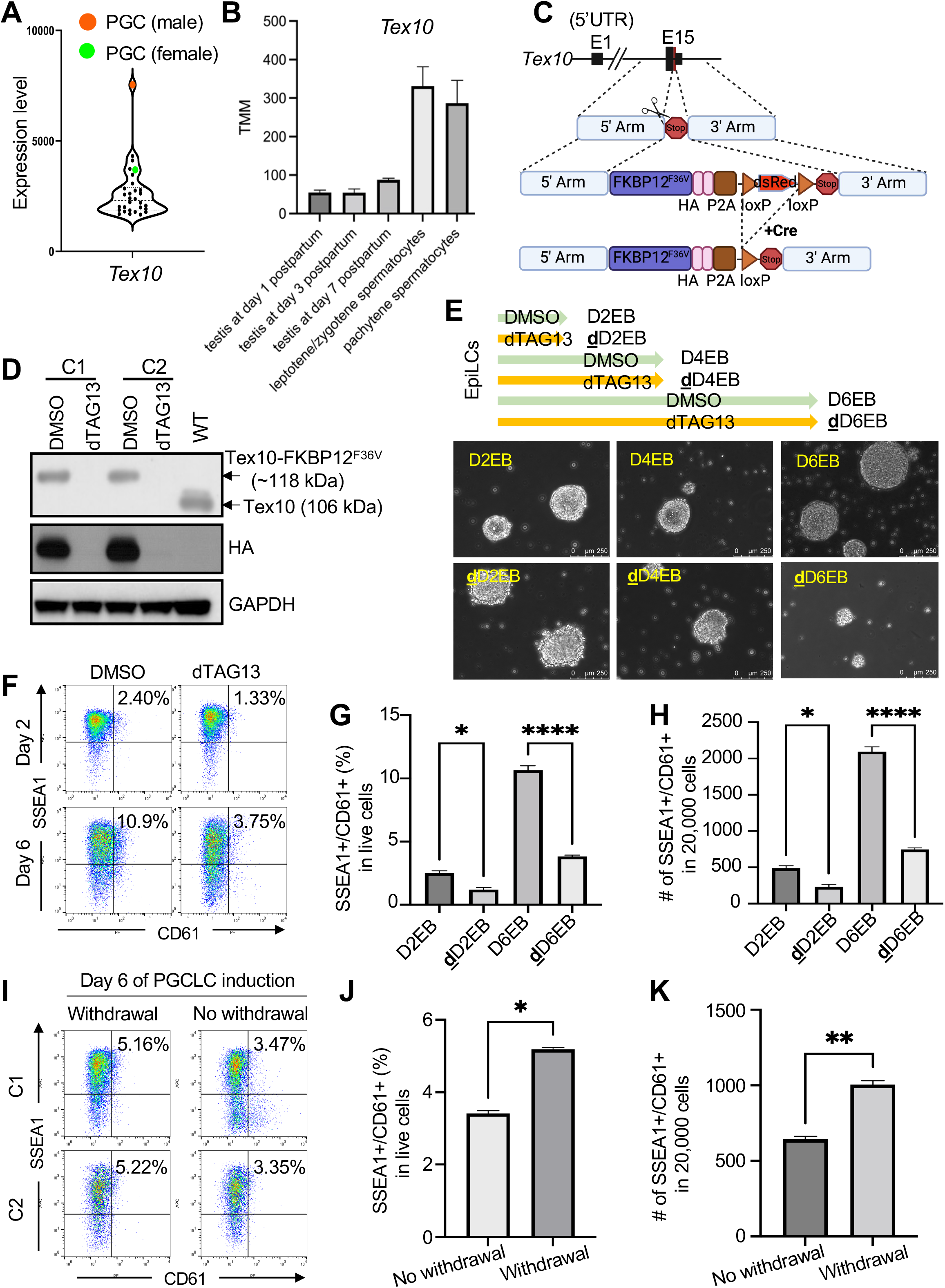
Loss of Tex10 compromises PGCLC specification efficiency. **(A)** Relative expression levels of Tex10 in mouse tissue/cell line samples (n=30). The male and female PGCs are indicated in orange and green, respectively. Data were obtained from the publicly available dataset (Hutchins et al. 2017). **(B)** Relative expression levels of *Tex10* on day 1, day 3, day 7 testes, and spermatocytes from adult testes. The data were curated from GSE83264. **(C)** Schematic representation of the Tex10 degron system. Briefly, the *Tex10* C terminus was edited using the CRISPR/Cas9 scissor to integrate degron system elements, and the Cre-loxP system was further used to delete the dsRed element. **(D)** Western blot validation on two cell clones (C1 and C2) of the Tex10 degron system. The top arrow indicates Tex10-HA-tagged FKBP12^F36V^ protein with a size of around 118 kDa, and the second arrow indicates the wildtype Tex10 protein of 106 kDa. **(E)** Cellular morphology of embryoid bodies (EBs) treated with DMSO or dTAG13 during six days of *in vitro* PGCLC specification from EpiLCs. **(F)** Flow cytometry analysis of PGCLC specification efficiency using cell surface markers SSEA1 and CD61. Percentages of double-positive (SSEA1^+^ and CD61^+^) cells are indicated at day 2 and day 6 of PGCLC induction for clone C1. **(G-H)** Quantification of double-positive (SSEA1^+^ and CD61^+^) percentage in live cells (G) and cell numbers per 20,000 analyzed cells (H) shown with bar plots. Two cell clones C1 and C2 were used as biological replicates, and an ANOVA test was used to detect significance. **(I-K)** Flow cytometry analysis of PGCLC specification efficiency using cell surface markers SSEA1 and CD61. Percentages of double-positive (SSEA1^+^ and CD61^+^) cells are indicated at day 6 of PGCLC induction (I). Quantification of double-positive percentage in live cells (J) and cell numbers per 20,000 analyzed cells (K) are shown with bar plots. Two cell clones C1 and C2 were used as biological replicates, and a paired t-test was used to detect significance. Withdrawal for depleting Tex10 only at day one of PGCLC induction and then transferring EBs into medium without dTAG13 to restore Tex10 expression; No Withdrawal for depleting Tex10 for six days of PGCLC induction.

### Tex10 depletion compromises the efficiency of PGCLC specification

We employed the well-established *in vitro* ESC-EpiLC-PGCLC differentiation system (Hayashi et al. 2011) with minor modifications (see Methods for details) to interrogate Tex10 functions in PGC development. Due to its required roles in the maintenance of ESCs and thus the inaccessibility to Tex10-deficient ESCs (Ding et al. 2015), we took advantage of the degron system (Nabet et al. 2018) (Fig. 1C) to deplete the Tex10 protein at specific periods during *in vitro* ESC-EpiLC-PGCLC differentiation. Using CRISPR targeting (Fig. 1C and see Methods for detail), we obtained and validated two Tex10 degron ESC clones (C1 and C2 in Fig. 1D). The Tex10 locus in these two clones was knocked in with the C-terminal in-frame HA-tagged FKBP12^F36V^ expression construct (Fig. 1C), resulting in a protein size of around 12 kDa larger than the wildtype (WT) Tex10 protein (Fig. 1D). Upon dTAG13 treatment, Tex10 is completely degraded within six hours (Fig. 1D). Following the *in vitro* PGC specification protocol (Hayashi et al. 2011), we compared the cellular morphology of cells treated with dTAG13 (Tex10 depletion) and cells treated with DMSO (control) for two, four, and six days. We found that, while the control cell clusters exhibited typical smooth boundaries of embryoid bodies (EBs) (Fig. 1E, D2/4/6EBs for embryoid bodies formed at the day2/4/6 PGCLC stages), the boundaries of dTAG13-treated <u>D</u>ay <u>2/4/6 EBs</u> (dD2/4/6EBs) became non-smooth and dispersed into small patches (Fig. 1E). We then performed DMSO/dTAG13 treatment during the initial ESC-to-EpiLC (24 and 48 h) differentiation and EB formation without cytokines used in the PGCLC medium and found that Tex10 depletion impairs EpiLC differentiation as well as EB formation (Fig. S1D and S1E). We further performed flow cytometry with SSEA1 and CD61, two typical PGC surface makers, and found that the number and ratio of double-positive cells, namely PGCLCs, were significantly decreased at day 2 and day 6 upon Tex10 depletion (Fig. 1F-H), consistent with the reduced expression of Prdm1 (Fig. S1F and S1G). Furthermore, we depleted Tex10 only on day 1 of PGCLC induction and then transferred EBs to a medium without dTAG13 to restore Tex10 expression. Compared to the group without dTAG13 withdrawal, the withdrawal group shows higher percentages of PGCLCs in live cells and more PGCLC numbers (Fig. 1I-K). These results support a critical role of Tex10 in PGCLC specification in addition to its function in cell survival.

### Germ cell development is dysregulated by Tex10 depletion at the PGCLC stage

To understand how Tex10 promotes the PGCLC specification, we investigated the transcriptional change induced by Tex10 depletion (Table S4). A total of 1831 genes were upregulated in D2EBs vs. EpiLCs, among which 282 genes were downregulated by Tex10 depletion (Fig. 2A). Gene ontology (GO) analysis indicated that “stem cell population maintenance” and “germ cell development” were disrupted, manifested by the downregulation of the pluripotency genes such as *Klf4* and *Tcl1*, and germ cell development genes, including *Prdm1, Prdm14*, and *Kit*, as well as *Esrrb* and *Nanog* with dual functions in pluripotency and germ cell development (Fig. 2A and S2A). On the other hand, 1278 genes were downregulated in D2EBs vs. EpiLCs, among which 326 genes were upregulated by Tex10 depletion. These genes are enriched in the GO terms such as “cell adhesion” and “positive regulation of apoptotic process” (Fig. 2A and S2A). To examine the overall early effects of Tex10 depletion on the expression of pluripotency, PGC, and lineage differentiation genes, we conducted the activity analysis (see Methods for details) on RNA-seq datasets of <u>**d**</u>D2EBs vs. D2EBs. We found that pluripotency and PGC regulation activities decreased, while mesoderm development activity increased after Tex10 depletion (Fig. 2B and S2B). More importantly, the PGC-restricting factor Otx2 (Zhang et al. 2018a) was also increased upon Tex10 depletion (Fig. 2C).

**Fig. 2.**
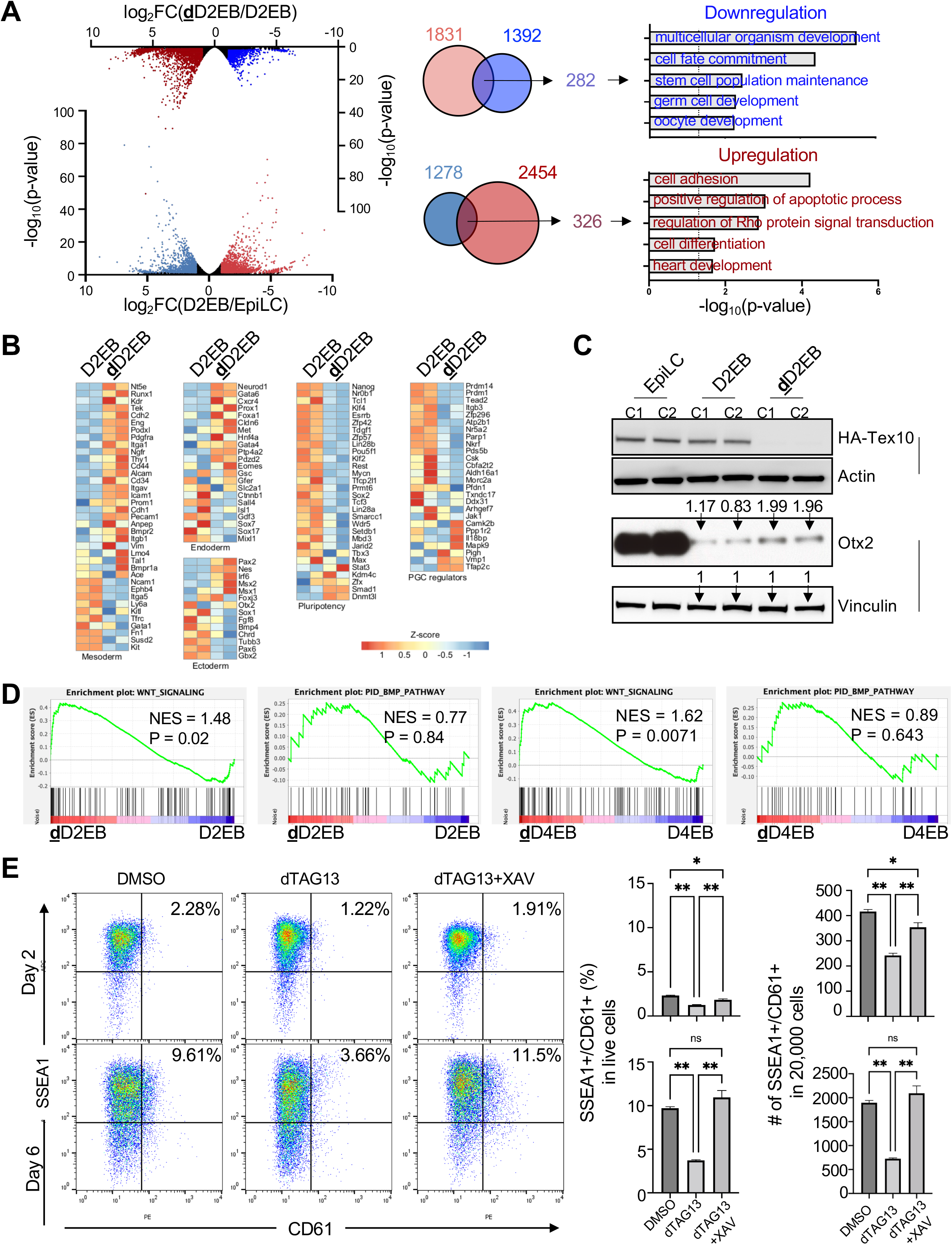
Wnt signaling is activated by Tex10 depletion to compromise the PGCLC specification. **(A)** Volcano plots show the comparison of transcriptomic profiles of day 2 PGCLC (D2EB) vs. EpiLC stage (bottom volcano), and dTAG13-treated day 2 PGCLC (dD2EB) vs. control (D2EB; top volcano). Venn diagrams show the intersections between 1831 genes upregulated in D2EB vs. EpiLC (log_2_FC > 1 & p-value < 0.05) and 1392 genes downregulated in dD2EB vs. D2EB (log_2_FC < −1 & p-value < 0.05), and between 1278 genes downregulated in D2EB vs. EpiLC (log_2_FC < −1 & p-value < 0.05) and 2454 genes upregulated in dD2EB vs. D2EB (log_2_FC > 1 & p-value < 0.05). Enriched gene ontology biological processes are shown for the two overlapping gene sets. **(B)** Heatmaps for gene expression profiles of pluripotency, PGC, ectoderm, mesoderm, and endoderm regulation markers in D2EBs vs. dD2EBs. **(C)** Western blots of key PGC markers in EpiLCs, D2EBs, and dD2EBs. Data from two Tex10-degron clones (C1 & C2) are shown with dTAG13 added at the EpiLC stage and Tex10 depletion happened in dD2EBs. Quantification values for Otx2 relative to Vinculin are shown on top of bands for D2 and dD2 EB samples. **(D)** Gene set enrichment analysis showing that Wnt signaling but not Bmp signaling is activated by Tex10 depletion on day 2 and day 4 of PGCLC induction. **(E)** Flow cytometry analysis of PGCLC specification efficiency using cell surface markers SSEA1 and CD61. Percentages of double positive (SSEA1^+^ and CD61^+^) cells are indicated at day 2 and day 6 of PGCLC induction for clone C1. Quantification of double-positive percentages in live cells and numbers per 20,000 analyzed cells are shown with bar plots. Two cell clones C1 and C2 were used as biological replicates, and an ANOVA test was used to detect significance.

To examine effects of Tex10 depletion at a different PGCLC stage, we treated the cells with dTAG13 for four days, resulting in 2494 (log_2_FC > 1 and p-value < 0.05) and 2454 (log_2_FC < −1 and p-value < 0.05) significantly upregulated and downregulated genes, respectively (Fig. S2C). Compared to two-day treatment, four-day treatment also compromised pluripotency and PGC specification, while lineage differentiation, particularly mesoderm development, was enhanced (Fig. S2D and S2E). Prdm1, Prdm14, and AP2γ are known to form the core PGC circuitry to initiate PGC-specific fate and repress somatic genes (Magnusdottir et al. 2013). However, we found that *Prdm1* and *Prdm14* were significantly downregulated upon Tex10 depletion (Fig. S2F top panels). These two factors also activate pluripotency genes (Tang et al. 2016). Accordingly, among the 162 genes consistently downregulated upon two-day and four-day dTAG13 treatments, genes with functional GO terms such as “stem cell population maintenance” and “germ cell and oocyte development” were significantly enriched (Fig. S2F). On the other hand, 172 genes were consistently upregulated and significantly enriched with functional GO terms on “apoptosis”, “cell adhesion”, and “heart development” in those two-day and four-day dTAG13-treated cells (Fig. S2F bottom panels).

Taken together, our data demonstrate that PGC development is positively regulated by Tex10, whose depletion leads to the disruption of the core PGC circuitry with a concomitant increase of apoptosis and somatic lineage development.

### Wnt signaling is hyperactivated by Tex10 depletion at the PGCLC stage

Wnt and Bmp signaling pathways are well recognized for their roles in regulating the PGC circuitry (Tang et al. 2016). Given that the core PGC circuitry is disrupted by Tex10 depletion, we asked whether these signaling pathways are also subject to Tex10 regulation. Gene set enrichment analysis (GSEA) of the Tex10 transcriptome revealed that the Wnt signaling pathway (NES = 1.48, p-value = 0.02), but not the Bmp signaling pathway (NES = 0.77, p-value = 0.84), was activated by Tex10 depletion at the early PGCLC stage (Fig. 2D left two panels). Such aberrant Wnt activation (i.e., hyperactivation) without a significant effect on Bmp signaling was also observed in <u>**d**</u>D4EBs vs. D4EBs (Fig. 2D right two panels). We further analyzed the expression of *Tex10, Prdm1, Prdm14*, and *Tfap2c* in ESCs, EpiLCs, PGCLCs, and EpiSCs based on the published microarray dataset (Hayashi et al. 2011). The expression levels of all these four factors were increased during the EpiLC to PGCLC stages and reached the highest in PGCLCs compared to ESCs, EpiLCs, and EpiSCs (Fig. S2G). Co-upregulation of *Tex10* with known PGC specifier genes *Prdm1, Prdm14*, and *Tfap2c* supports its potential role in PGC specification by controlling Wnt activity. Remarkably, the Wnt signaling positive regulators, such as *Wnt4* and *Wnt6*, were upregulated, whereas the Wnt signaling negative regulators, such as *Grb10* (Tezuka et al. 2007), *Sfrp1* (Rattner et al. 1997), *Psmd3* (Aberle et al. 1997; Kominami et al. 1997; Fararjeh et al. 2019), and *Psmd7* (Yi et al. 2017), were downregulated upon Tex10 depletion at both day 2 and 4 PGCLC stages (Fig. S2H). Moreover, the Wnt inhibitor XAV939 (10 μM, the same concentration used in (Hackett et al. 2018)) rescued the Tex10 depletion phenotypes by increasing the percentages of PGCLCs in live cells and PGCLC numbers (Fig. 2E and S2I). In PGC specification, the Bmp4-induced Wnt signaling pathway activates *T* and inhibits *Otx2*, leading to the activation of *Prdm1* and *Prdm14* (Yao et al. 2022). *T* is, however, not identified in our initial RNA-seq (Fig. 2A-B), possibly due to insufficient reads, because it is indeed upregulated when we collected more EBs and repeated RNA-seq, although this *T* increase upon Tex10 depletion is not as significant as *Otx2* upregulation (Fig. S2J). Therefore, depletion of Tex10 may compromise PGC development mainly through aberrant activation of both Wnt signaling and the PGC restriction factor Otx2 during the specification stage, distinct from the classic effects of Bmp4-induced Wnt signaling on PGC development (see Discussion).

To test whether Tex10 directly targets and regulates the Wnt signaling regulatory genes, we performed ChIP-seq for Tex10 and H3K4me3 in D2EBs. We identified 4331 peaks (Table S5) at the D2PGCLC stage, out of which ∼70% peaks (3015/4331, corresponding to 2411 nearest genes) co-existed with the H3K4me3 histone mark (Table S6), indicating that Tex10 bound to the potential promoter regions (Fig. 3A). We found 322 common genes after intersecting Tex10-bound/H3K4me3-marked genes (n = 2411) with genes that were downregulated upon Tex10 depletion (n = 3076, log_2_FC < −0.5 and p-value < 0.05) (Fig. 3B and Table S7). GO analysis revealed that these genes were enriched with functional GO terms such as “negative regulation of apoptosis” (e.g., *Trim27* and *Tyro3*) and “negative regulation of Wnt signaling” (e.g., *Psmd3* and *Psmd7*) (Fig. 3C, 3D and Table S8). On the other hand, we found 252 genes (Table S9) after intersecting Tex10-bound/H3K4me3-marked genes with genes that are upregulated upon Tex10 depletion (n = 3241, log_2_FC > 0.5 and p-value < 0.05, Table S1). The highest enriched GO terms were related to cell proliferation, microvillus assembly, and negative regulation of TOR signaling (Fig. S3A, S3B, and Table S10); however, no GO terms related to Wnt signaling were enriched in the above 252 genes (Table S10, Fig. S3A, and S3C). We further validated the direct binding of Tex10 to the promoters of *Psmd3* and *Psmd7* (Fig. 3E), which are components of the 19S proteasome subcomplex with reported roles in finetuning the Wnt/beta-catenin signaling (Yi et al. 2017). In particular, PSMD7 is directly associated with beta-catenin degradation from the proteasome (Yi et al. 2017). We further knocked down Psmd7 using two shRNAs (Fig. S3D and S3E) and showed that Psmd7 knockdown compromised PGCLC specification efficiency (Fig. 3F), thus mimicking the Tex10 depletion phenotypes. In contrast, the ectopic expression of Psmd7 partially rescued the PGCLC specification efficiency of Tex10-depleted (i.e., dTAG13-treated) cells (Fig. 3G, S3F, and S3G). Thus, these results indicate that Tex10 directly activate the transcription of Wnt signaling negative regulators and support that Tex10 depletion compromises the PGCLC specification by aberrant activation of Wnt signaling.

**Fig. 3.**
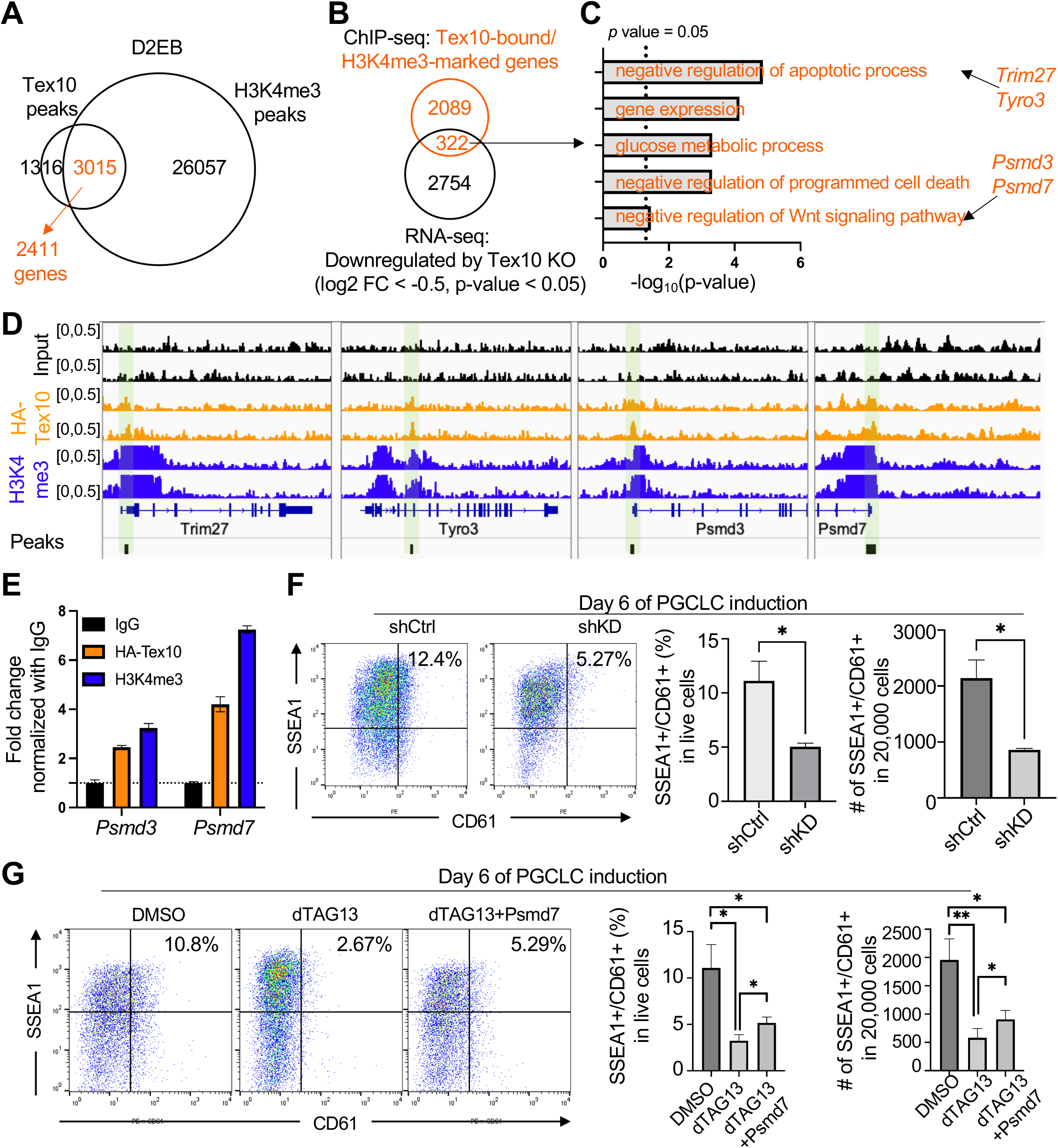
Wnt negative regulators Psmd3/7 are targeted and downregulated by Tex10 depletion to compromise PGCLC specification. **(A)** Venn diagram showing the overlap between Tex10 and H3K4me3 ChIP-seq peaks at the D2PGCLC stage. **(B)** Venn diagram showing the intersection between Tex10-bound/H3K4me3-marked genes and genes downregulated upon Tex10 depletion. **(C)** Gene ontology biological processes enriched in the 322 genes shown in B. **(D)** Genome browser tracks of Tex10 and H3K4me3 ChIP-seq signals near *Trim27, Tyro3, Psmd3*, and *Psmd7* gene loci in D2EBs. Green shaded regions indicate the Tex10 peaks. **(E)** ChIP-qPCR validation of Tex10 binding and H3K4me3 mark at the promoter regions of *Psmd3* and *Psmd7*. Tex10 ChIP experiment was performed with an anti-HA tag antibody, and IgG served as a negative control.” **(F)** Flow cytometry analysis of PGCLC specification efficiency using cell surface markers SSEA1 and CD61. Percentages of double-positive (SSEA1^+^ and CD61^+^) cells are indicated at day 6 of PGCLC induction for control shRNA (shCtrl) and Psmd7 shRNA (shKD). Quantification of double-positive percentages in live cells and numbers per 20,000 analyzed cells are shown with bar plots. N = 2 biological replicates per condition (two control shRNAs vs. two Psmd7 shRNAs), and an unpaired t-test was used to detect significance. **(G)** Flow cytometry analysis of PGC specification efficiency using the cell surface markers SSEA1 and CD61. Percentages of double-positive (SSEA1^+^ and CD61^+^) cells are indicated at day 6 of PGCLC induction for DMSO, dTAG13, and dTAG13 plus Psmd7 ectopic expression (dTAG13+Psmd7) treatment. Quantification of double-positive percentages in live cells and numbers per 20,000 analyzed cells are shown with bar plots. Two cell clones C1 and C2 were used as biological replicates, and a paired t-test was used to detect significance.

### Tex10 overexpression enhances PGC specification efficiency

Given that Tex10 interacts with and regulates Nanog in mouse ESCs (Ding et al. 2015) and that Nanog acts as a potent inducer of PGC specification from ESC-derived EpiLCs (Murakami et al. 2016), we next asked if Tex10 overexpression would promote PGC development. We first established two ESC clones transgenic for FLAG-tagged Tex10 and confirmed Tex10 overexpression (OE) relative to empty-vector (EV) control cells (see Methods for details and Fig. S4A). We then conducted the *in vitro* PGC specification to compare cellular morphology at different stages between OE and EV control ESCs. Our results showed that the cellular morphology was similar (Fig. 4A and S4B) and that Tex10 OE significantly increased the efficiency measured by SSEA1 and CD61 dual positivity (Fig. 4B and S4C). We also noted that, while both RNA and protein levels of Tex10 were increased throughout the entire process, Nanog and Otx2 became appreciably upregulated and downregulated, respectively, at the late PGCLC stage (D6EB, Fig. 4C and 4D). Finally, to investigate whether Tex10 OE is sufficient to induce PGCLCs without cytokines, we treated EV and OE cells with (+) or without (-) cytokines (see Methods for cytokines information). Our results showed that Tex10 OE was not sufficient to induce PGCLCs without cytokines (Fig. 4E-G and S4D). These data further confirm the role of Tex10 in promoting *in vitro* PGC specification under the PGCLC culture condition.

**Fig. 4.**
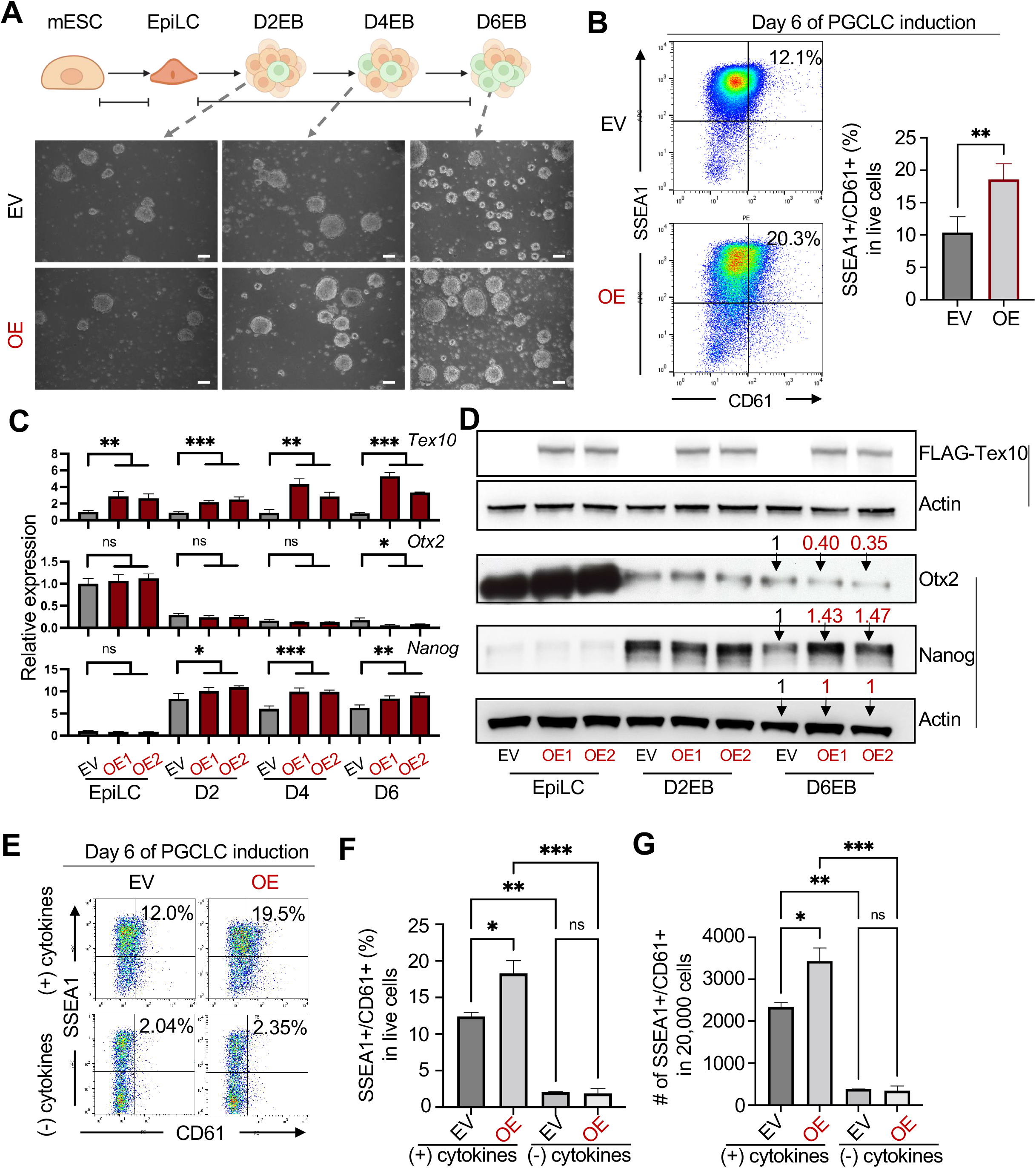
Tex10 overexpression enhances PGCLC specification efficiency. **(A)** Cellular morphology of Tex10 overexpression (OE) vs. control EBs (Empty vector, EV) at D2, D4, and D6 PGCLC stages (scale bar, 100 μm). **(B)** Flow cytometry analysis of the efficiency of PGCLC specification using the markers SSEA1 and CD61. Percentages of double-positive (SSEA1+ and CD61+) cells are indicated at day 6 of PGCLC induction. Two cell clones, OE1 and OE2, were used as biological replicates and the paired t-test was used to detect significance. **(C)** Quantitative RT-PCR analysis of *Tex10, Otx2*, and *Nanog* expression during the transition from EpiLCs to D2, D4, and D6 EBs. ANOVA was performed for the significance test. **(D)** Western blot analysis of FLAG-Tex10, Otx2, and Nanog expression during the transition from EpiLCs to D2, D4, and D6 EBs. Quantification values for Otx2 and Nanog relative to Actin are shown on the top bands for D6EB samples. **(E-G)** Flow cytometry analysis of PGCLC specification efficiency using cell surface markers SSEA1 and CD61. Percentages of double-positive (SSEA1+ and CD61+) cells are indicated at day 6 of PGCLC induction with (+) or without (-) cytokines (E). Quantification of double-positive percentages in live cells (F) and numbers per 20,000 analyzed cells (G) are shown with bar plots. Two cell clones, OE1 and OE2, were used as biological replicates. The ANOVA test was used to detect significance.

### Tex10 overexpression restricts Wnt signaling and somatic lineage programs to promote pluripotency and PGC development

To gain molecular insight into Tex10 OE in promoting PGCLC specification, we investigated the time-course transcriptional changes during ESC-EpiLC-PGCLC differentiation (Table S11). We first confirmed the ectopic OE of *Tex10* at all stages (EpiLC, D2, D4, and D6 PGCLC stages, Fig. 5A). Tex10 OE increased PGCLC specification markers (*Dazl* and *Kit*) and PGC regulator *Nr5a2* in D6EBs. The core PGC circuitry genes (*Prdm1, Prdm14*) and the PGC specification regulator *Zfp296* were appreciably (although not significantly) upregulated in D2EBs (for *Prdm1/14*) or throughout the time course (for *Zfp296*) upon Tex10 OE and maintained at relatively high expression levels throughout the PGCLC stages (D2, D4, D6 PGCLC stages) compared to the EpiLC stage (Fig. 5B).

**Fig. 5.**
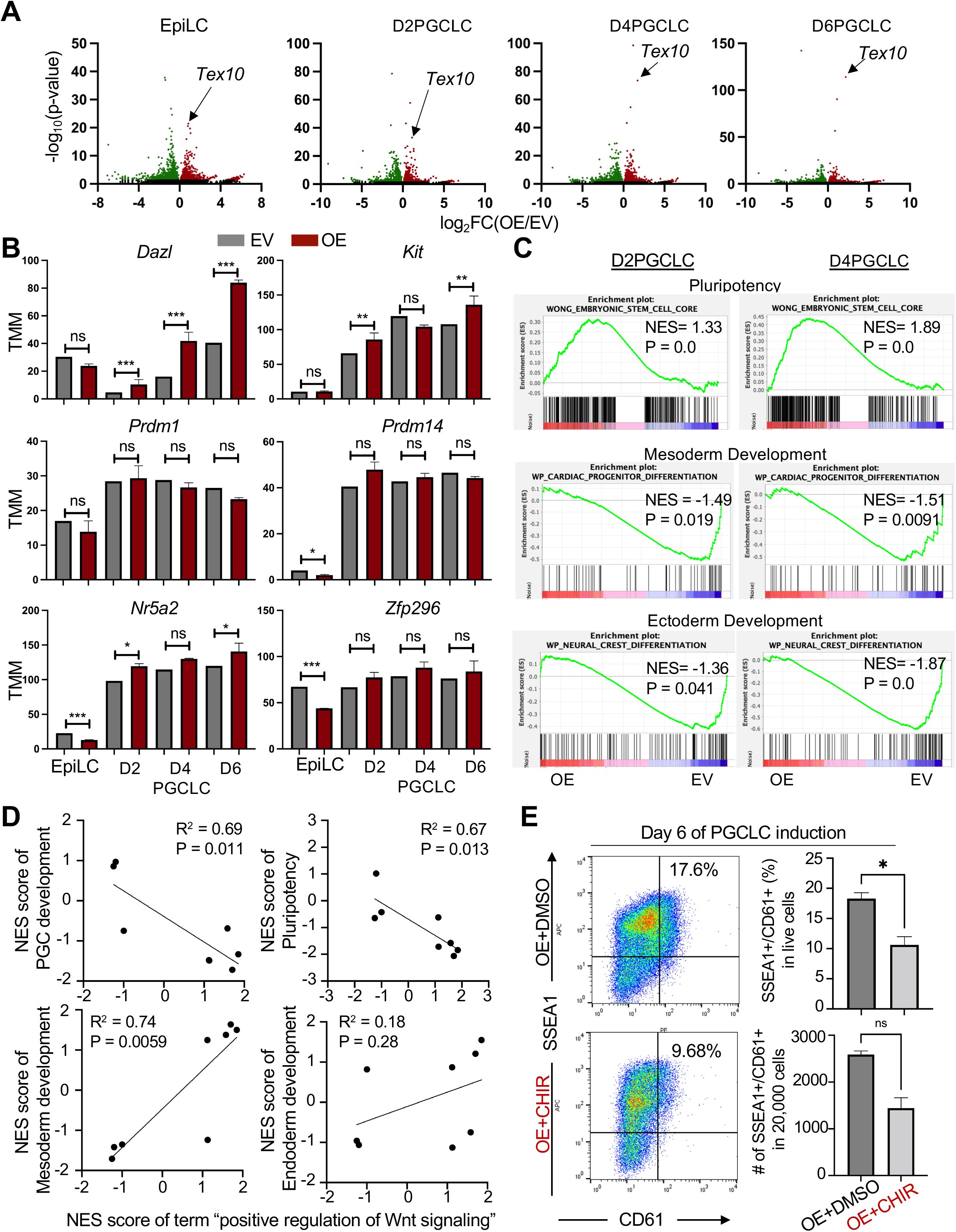
Tex10 overexpression restricts the Wnt signaling and somatic lineage programs in promoting pluripotency and PGC development. **(A)** Volcano plots depicting global transcriptomic changes induced by Tex10 OE at EpiLC, D2, D4, and D6 PGCLC stages. **(B)** Bar plots showing expression changes of representative PGC regulators induced by Tex10 OE during the transition from EpiLC to D6PGCLC stage. The p-value was computed by edgeR using a generalized linear model-based statistical method. **(C)** Gene set enrichment analysis of the effects of Tex10 OE on pluripotency, mesoderm, and ectoderm development at D2 and D4 PGCLC stages. **(D)** Regression models exploring the correlation between NESs (normalized enrichment scores) of the positive regulation of Wnt signaling and PGC development, pluripotency regulation, mesoderm, and endoderm development. **(E)** Flow cytometry analysis of the effect of the Wnt activator (CHIR: CHIR99021) on PGCLC specification efficiency in Tex10 OE cells. Percentages of double-positive (SSEA1+ and CD61+) cells are indicated at day 6 of PGCLC induction. Quantification of double-positive percentages in live cells and numbers per 20,000 analyzed cells are shown with bar plots. Two cell clones, OE1 and OE2, were used as biological replicates and the paired t-test was used to detect significance.

Considering the potential competition between PGC specification and somatic lineage differentiation (Ohinata et al. 2005; Kimura et al. 2014), we further used GSEA to interrogate the global gene expression changes upon Tex10 OE by focusing on somatic lineage differentiation and pluripotency genes. We found that pluripotency genes were significantly upregulated while mesoderm and ectoderm development genes were downregulated considerably upon Tex10 OE (Fig. 5C). Consistent with this, multilineage development genes were enriched in downregulated genes at the EpiLC (Fig. S5A), D2 (Fig. S5B), D4 (Fig. S5C), and D6 (Fig. S5D) PGCLC stages in Tex10 OE cells. On the other hand, the “stem cell population maintenance” GO term was enriched in upregulated genes at both D4 and D6 PGCLC stages (Fig. S5C and S5D). In addition, the GO term “DNA methylation” was also enriched in downregulated genes at EpiLC, D2, and D4 PGCLC stages (Fig. S5A-C), in line with our previous finding of Tex10 functions in DNA demethylation (Ding et al. 2015). More importantly, “Wnt signaling” (especially canonical Wnt signaling) related GO terms were enriched in downregulated genes at D2, D4, and D6 PGCLC stages in Tex10 OE cells (Fig. S5B-D). Moreover, the Tex10-targeted Wnt negative regulator *Psmd7* was upregulated by Tex10 OE throughout all PGCLC stages, and *Psmd3* was upregulated by Tex10 OE at both D4 and D6 PGCLC stages (Fig. S5E).

Through systematic investigation of publicly available RNA-seq data of known PGC regulators (Nr5a2, Hdac3, and Zfp296) and our Tex10 RNA-seq data in the context of Wnt signaling regulation and PGC development, we found that Tex10 depletion, *Nr5a2* KO (Hackett et al. 2018), and *Hdac3* knockdown (Mochizuki et al. 2018) datasets all showed the activation of Wnt signaling, whereas *Zfp296* KO (Hackett et al. 2018) and Tex10 OE showed the inhibition of Wnt signaling (Fig. S5F). Based on existing data and prior knowledge that both aberrant Wnt activation and inhibition impair PGC specification (Hackett et al. 2018), we created linear regression models to explore the correlation of the GO term “positive regulation of Wnt signaling” with PGC regulation, pluripotency regulation, and lineage development in the context of Tex10, Hdac3, and Nr5a2 regulation. We found that the GO term “positive regulation of Wnt signaling” is negatively correlated with PGC and pluripotency regulation (Fig. 5D) but positively associated with mesoderm development at the PGC specification stage (Fig. 5D). No apparent correlation was observed between Wnt signaling and endoderm development (Fig. 5D). This mathematical model supports that Tex10 OE attenuates Wnt signaling to promote PGC development and repress multilineage differentiation. However, the possible functional interactions between Tex10 and these factors remain unknown. To experimentally test the results from these linear regression models, we treated Tex10 OE cells with the Wnt activator CHIR99021 (3 μM, the same concentration used in (Hackett et al. 2018)). As predicted, activation of Wnt signaling compromised the efficiency of specification induced by Tex10 OE (Fig. 5E). Together, we conclude that Tex10 overexpression restricts the Wnt signaling and somatic lineage programs in promoting pluripotency and PGC development.

### Tex10 is functionally important for male fertility partly through the control of Wnt signaling

Tex10 conventional knockout embryos die at the preimplantation stage (Ding et al. 2015), precluding the analysis of its functional contribution to PGC development *in vivo*. Therefore, we employed CRISPR genome editing to create a conditional allele of Tex10 and obtained homozygous *Tex10* f/f mice (see Method for details, Fig. S6A). To investigate Tex10 functions in PGC development *in vivo*, we first mated *Tex10* f/f mice with *Prdm1*-Cre mice, which express Cre recombinase under the control of the early PGC marker *Prdm1* promoter (Ohinata et al. 2005), to obtain *Tex10* f/+;*Prdm1*-Cre/+ mice, which were then mated with *Tex10* f/+ mice (Fig. S6B). We first examined genotypes of liveborn pups from five litters, among which were 7 Group1 (genotypes of *Tex10* +/+, f/+, with or without Cre), 7 Group2 (genotypes of *Tex10* +/-with or without Cre, and *Tex10* f/-without Cre) pups, and surprisingly no *Tex10* f/-;*Prdm1*-Cre/+ (cKO) pups (Fig. S6B). Considering the low numbers of pups from each litter, we further examined 30 decidual swellings obtained at 12.5 *dpc* (*days post coitum*) from the same mating strategy. We found 19 blighted ova, 9 Group1 (genotypes of *Tex10* +/+, f/+, with or without Cre) embryos, 2 Group2 (genotypes of *Tex10* +/-;*Prdm1*-Cre/+ and *Tex10* f/-without Cre) embryos, and again no cKO embryos (Fig. S6C and S6D). The protein level of Tex10 is reduced to approximately 50% in HET (*Tex10* +/- and f/-without Cre) compared to WT (*Tex10* f/+, +/+ without Cre) E12.5 embryos (Fig. S6E), supporting the effectiveness of the conditional knockout strategy. The surprising lack of cKO embryos prevented us from investigating the exact effect of Tex10 depletion on the specification of PGC *in vivo* using *Prdm1*-Cre mice. It raised the question of how the PGC developmental defect induced by Tex10 deficiency would cause embryonic lethality at E12.5 (see Discussion).

On the other hand, the constitutive human WNT/CTNNB1 activation could trigger male germ cell depletion (Chassot et al. 2017). Tex10 is specifically and highly enriched in testis (Fig. 1B and S1C), and its depletion hyperactivates Wnt signaling, as indicated by our *in vitro* data. Therefore, we decided to directly investigate whether the germline quality is compromised in the Tex10 KO testes. First, we confirmed the expression of Tex10 in the testes using our previously constructed *Tex10* LacZ/+ mice that harbor a LacZ cassette at the *Tex10* locus (Ding et al. 2015). Beta-galactosidase (the coding product of the LacZ gene) was used to locate Tex10-expressing cells. We also used Sox9 to mark Sertoli cells (Hemendinger et al. 2002), the epithelial supporting cells of seminiferous tubules. Indeed, Tex10 was expressed in differentiating germ cells in seminiferous tubules (Fig. 6A). Then, we conditionally KO Tex10 using male germline-specific *Stra8*-iCre (Sadate-Ngatchou et al. 2008) acting during the early-stage spermatogonia to pre-leptotene-stage spermatocyte development (Fig. 6B). Tex10 cKO with *Stra8*-iCre mice had fewer sperms (Fig. 6C) and reduced sperm mobility (Video S1 for WT and S2 for cKO). More importantly, male *Tex10* f/f;*Stra8*-iCre mice showed compromised fertility compared to WT mice based on litter size (Fig. 6D). Although Tex10 is a “testis-specific transcript-10” for its namesake, Tex10 was surprisingly also expressed in female PGC (Fig. 1A, ranked#5) and the growing oocyte (Fig. S1B and S6F). Thus, we employed female germline-specific *Zp3*-Cre, acting in the growing oocyte before the completion of the first meiotic division (de Vries et al. 2000) to delete Tex10 in oocytes (Fig. S6G). We found no noticeable difference in ovary morphology or female fertility between WT and cKO mice using *Zp3*-Cre (Fig. S6H and S6I), supporting a preferred role of Tex10 in male germline development.

**Fig. 6.**
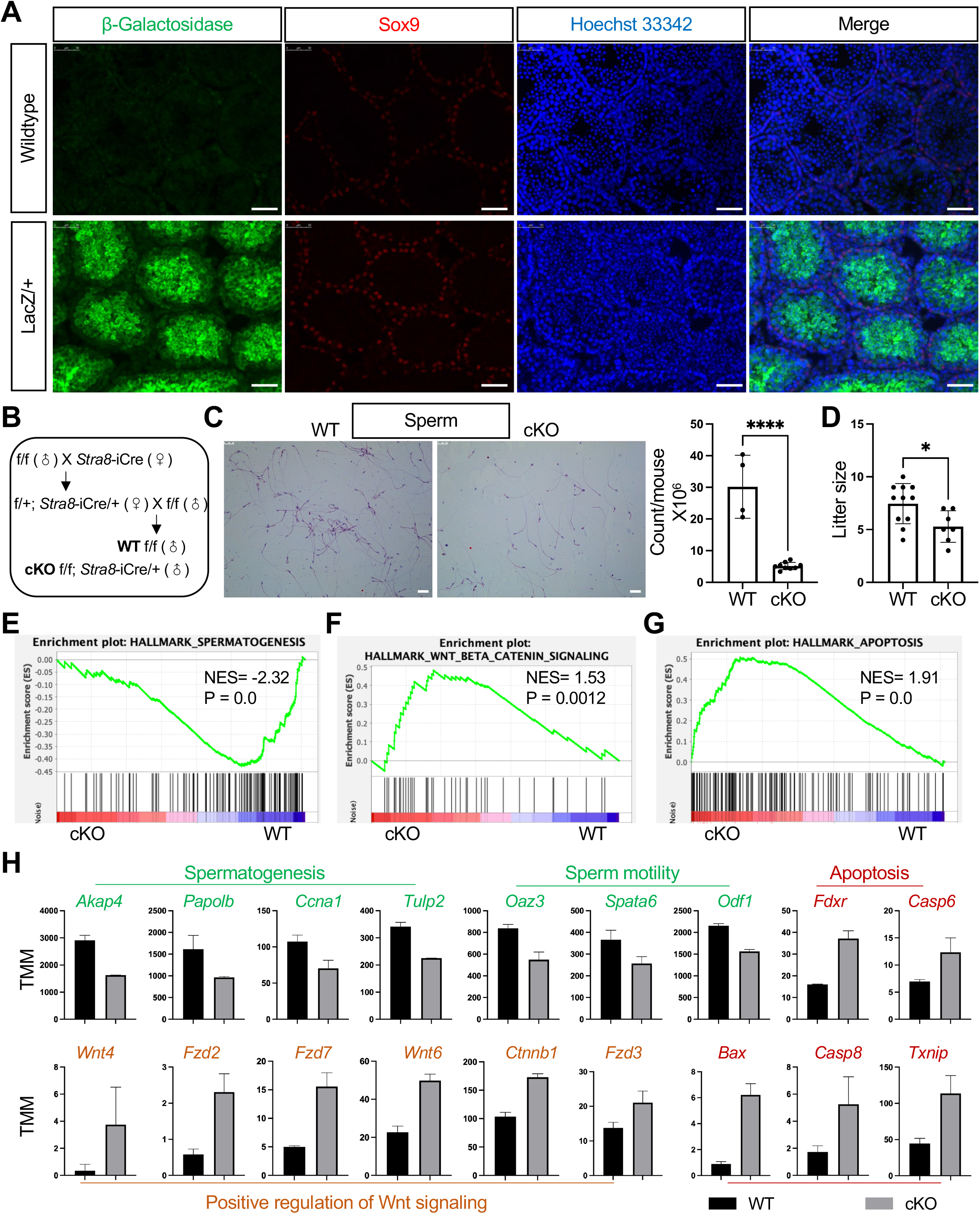
Tex10 cKO compromises male fertility. **(A)** Immunofluorescence images showing protein expression of β-Galactosidase (the coding product of the LacZ gene) and Sox9 in wildtype and *Tex10* LacZ/+ mouse testes. Hoechst 33342 shows the nucleus (scale bar, 50 μm). **(B)** Schematic depiction of the mating strategy for generating cKO (*Tex10* f/f;Stra8-iCre/+) and WT-equivalent (*Tex10* f/f) male mice. **(C)** Morphology and quantification of sperm count between cKO and WT male mice from the matings shown in B (scale bar, 25 μm). **(D)** Comparison of litter sizes produced by *Tex10* +/+ female mice mated with male WT and cKO mice from the matings shown in B. **(E-G)** GSEA plots of spermatogenesis (E), Wnt/β-catenin signaling (F), and apoptosis (G) obtained using RNA-seq data from testes of WT and Tex10 cKO mice from the matings shown in B. **(H)** Bar plots showing gene expression changes related to spermatogenesis, sperm motility, apoptosis, and Wnt signaling between WT and Tex10 cKO mice from matings shown in B.

Since spermatogenesis was defective upon Tex10 depletion, we extracted the RNA from the testes of cKO and WT mice with *Stra8*-iCre, followed by RNA sequencing to explore potential mechanisms. First, we confirmed that Tex10 was significantly downregulated in the testes of cKO mice (log_2_FC = −1.1, FDR = 8.23E-05, Table S12). GSEA on the hallmark dataset further showed that only one gene set, namely spermatogenesis, was significantly compromised upon Tex10 cKO (Fig. 6E). In contrast, the Wnt signaling and apoptosis were upregulated (Fig. 6F and 6G). In particular, spermatogenesis genes, such as *Akap4* (Fang et al. 2019), *Papolb* (Kashiwabara et al. 2002; Zhuang et al. 2004), *Ccna1* (Liu et al. 1998; Liu et al. 2000; Salazar et al. 2005), and *Tulp2* (Zheng et al. 2021), and sperm motility related genes, such as *Oaz3* (Tokuhiro et al. 2009), *Spata6* (Yuan et al. 2015), and *Odf1* (Fawcett and Phillips 1969; Woolley 2010; Yang et al. 2012; Chen et al. 2016), were all downregulated; whereas apoptosis (e.g., *Bax, Casp6*) and Wnt signaling genes (e.g., *Fzd2, Ctnnb1*) were upregulated (Fig. 6H). We further confirmed that *Tex10, Psmd3*, and *Psmd7* were downregulated in *Tex10* f/-;*Stra8*-iCre/+ (i.e., Tex10 cKO) mouse testis compared to WT (i.e., *Tex10* f/f) testis (Fig. S6J). Furthermore, we curated genes upregulated by β-catenin accumulation from the Molecular Signatures Database (gene set ID: M5895, (Subramanian et al. 2005; Liberzon et al. 2015)) and found that this gene set is appreciably upregulated by Tex10 KO *in vivo* and *in vitro* (Fig. S6K), consistent with our previous findings (Fig. 2D and 6F). Taken together, these results partly explain the abnormal sperm phenotypes and lend further support for the roles of Tex10 in proper male germline development through regulating the Wnt signaling.

### Tex10 is critical for round spermatic formation during spermatogenesis through the control of Wnt signaling

To further investigate which stage of spermatogenesis is compromised by Tex10 depletion, we collected cells from the testes of WT (*Tex10* +/+;*Stra8*-iCre/+) and cKO (*Tex10* f/-;*Stra8*-iCre/+) mice. We first employed the FACS approach as described (Gaysinskaya et al. 2014) to distinguish cells from different stages (Fig. 7A-B). Although we could not perfectly distinguish all different populations, we identified one round <u>s</u>permatid (RS)-like and one “<u>s</u>permato<u>g</u>onia and <u>s</u>permato<u>c</u>yte (Sg_Sct)-like” cell populations (Fig. 7B left panels). Importantly, we found that Tex10 depletion significantly decreased the RS-like population. In contrast, the Sg_Sct-like population was not significantly affected (Fig. 7B right two bar charts). Supporting this finding, the RNA-seq data from whole testes showed that 14 out of 15 round spermatid marker genes (Hermann et al. 2018) were significantly downregulated by Tex10 depletion (Fig. 7C). We also checked three spermatogonia marker genes (i.e., *Nanos3, Sohlh1* and *Sohlh*2) and four spermatocyte marker genes (i.e., *Sycp1, Sycp3, Piwil1 and Piwil2*) genes. We found a non-significant effect on both sets of genes, except for *Sohlh2* (Fig. S7A).

**Fig. 7.**
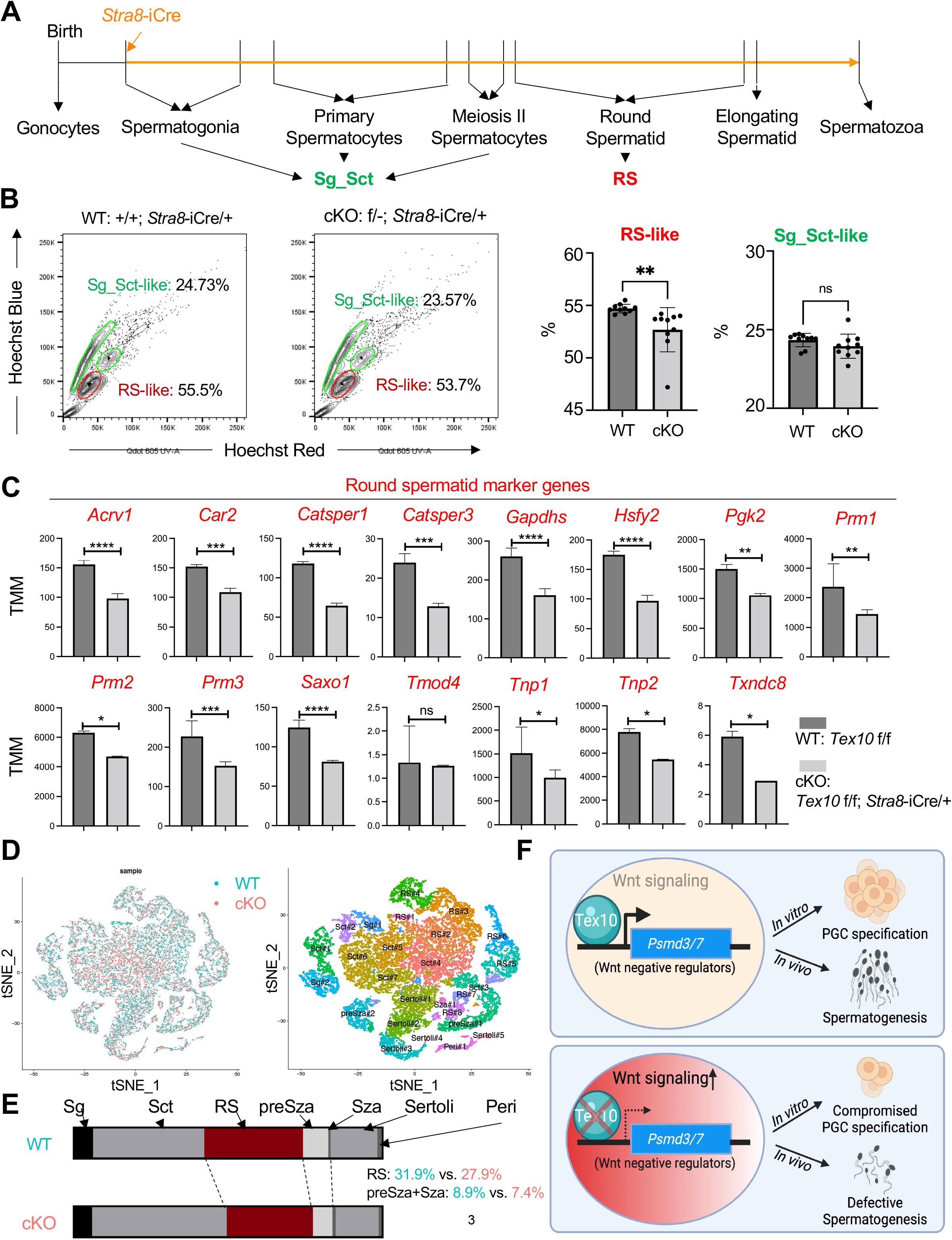
Round spermatid formation is compromised in Tex10 cKO mice. **(A)** Schematic representation of mouse spermatogenesis. An orange arrow indicates the stage for *Stra8*-iCre to play effect. Sg for spermatogonia, Sct for spermatocyte, and RS for round spermatid. **(B)** Representative flow cytometry isolation of spermatogenic populations in mouse testes (left) and quantification data (right). N = 10 biological replicates per condition, and an unpaired t-test was used to detect significance. **(C)** Expression changes of round spermatid markers in WT (*Tex10* f/f) vs. cKO (*Tex10* f/f;*Stra8*-iCre/+) mouse testes. The p-value was computed by edgeR using a generalized linear model-based statistical method. **(D)** The scRNA-seq data are colored by WT (*Tex10* +/+;*Stra8*-iCre/+, left panel, blue) and cKO (*Tex10* f/-;*Stra8*-iCre/+, left panel, pink), and the scRNA-seq identifies cell populations from mouse testis (right panel), and n = 2 biological replicates per condition. t-SNE, t-distributed stochastic neighbor embedding. **(E)** Cell proportion changes in WT (*Tex10* +/+;*Stra8*-iCre/+) vs. cKO (*Tex10* f/-;*Stra8*-iCre/+) mouse testes. Sg for spermatogonia, Sct for spermatocyte, RS for round spermatid, preSza for pre spermatozoa, Sza for spermatozoa, Peri for peritubular, and Sertoli for Sertoli cells. **(F)** A summary model of Tex10 in controlling PGC specification and spermatogenesis. Under the wildtype condition, Tex10 binds the promoters and upregulates the expression of *Psmd3/7*, negative regulators for Wnt signaling, leading to the attenuation of the Wnt pathway (upper panel). Upon Tex10 depletion, the downregulation of *Psmd3/7* expression levels hyperactivates Wnt signaling, resulting in compromised PGCLC specification *in vitro* and spermatogenesis *in vivo* (bottom panel).

To gain further insight into the mechanisms underlying Tex10 regulation of spermatogenesis, we conducted single-cell RNA-sequencing on WT (*Tex10* +/+;*Stra8*-iCre/+) and cKO (*Tex10* f/-;*Stra8*-iCre/+) testis (Fig. 7D), yielding well-defined different cell populations clustered by specific markers (Fig. S7B). Consistent with our findings described above, we confirmed that the RS population was decreased in cKO relative to WT (Fig. 7E). Among different RS clusters, RS#2 and RS#3 showed the most obvious decrease (Fig. S7C), and RS markers *Tnp1* and *Tnp2* were significantly decreased in both RS#2 and RS#3 of cKO relative to WT samples (Fig. S7D). In particular, the GO term “male meiosis I” was enriched for upregulated genes in RS#3 of cKO relative to WT, indicating that spermatogenesis might be arrested at the meiosis I stage (Fig. S7E). Moreover, the GO term “positive regulation of Wnt signaling” was also significantly enriched (p-value < 0.05) for upregulated genes in RS#3 of cKO relative to WT, supporting Wnt signaling activation by Tex10 depletion during spermatogenesis, especially the round spermatid formation (Fig. S7E). These results strongly support the functions of Tex10 in regulating round spermatid formation of spermatogenesis, partially through regulating Wnt signaling, a mechanistic connection previously reported in spermatogonia regulation and consequently spermatogenesis (Takase and Nusse 2016; Tokue et al. 2017).

Interestingly, we found that patients with severely impaired spermatogenesis show significant downregulation of *TEX10* (Fig. S7F and Table S13), which is also observed in infertile teratozoospermia individuals compared with fertile normospermia individuals (Fig. S7G and Table S14), suggesting that TEX10 may also be important for male fertility in humans. As constitutive human WNT/CTNNB1 activation could trigger male germ cell depletion (Chassot et al. 2017), Tex10 functions in finetuning the Wnt signaling in PGC and sperm development are likely evolutionarily conserved.

## DISCUSSION

Identifying an entire repertoire of PGC regulatory network genes will be instrumental in unraveling the molecular mechanisms underlying mammalian germ cell development. CRISPR screen has been applied to *in vitro* PGC specification leading to the identification of additional PGC specification regulators such as Zfp296 and Nr5a2 (Hackett et al. 2018). In this study, we created first-of-its-kind Tex10-degron ESC system and Tex10 conditional knockout mouse model to overcome its preimplantation requirements (Ding et al. 2015), which enabled us to discover Tex10 as a previously unappreciated player in PGC specification *in vitro* and male germline quality control *in vivo*. Mechanistically, we found that Tex10 directly binds and transcriptionally activates Wnt negative regulators such as Psmd3/7 in maintaining the precise control of Wnt signaling for proper PGCLC specification and spermatogenesis. However, upon Tex10 depletion, the Wnt signaling is misregulated (i.e., hyperactivated) due to the downregulation of those negative regulators, resulting in compromised PGCLC specification and defective spermatogenesis with direct relevance to human patients (Fig. 7F, Fig. S7F and S7G). On the other hand, Tex10 overexpression induces Wnt repression (Fig. S5B-D) and restricts somatic lineage programs (Fig. 5C), leading to increased PGCLC specification. Taken together, our study further expands the PGC regulatory network and provides a potential therapeutic target for treating male infertility.

Pluripotency reactivation is a notable feature of the PGC specification (Magnusdottir et al. 2013; Tang et al. 2016). For the core pluripotency genes Nanog, Oct4, and Sox2 (Boyer et al. 2005; Festuccia et al. 2013), conditional deletion of either Sox2 (Campolo et al. 2013) or Oct4 (Kehler et al. 2004) results in PGC death, and the induced knockdown of Nanog also results in PGC apoptosis (Yamaguchi et al. 2009). In addition, the deletion of the Nanog target gene Esrrb reduces PGC numbers *in vivo* (Mitsunaga et al. 2004). Along this line, we found that depletion of Tex10, an interaction partner of Nanog, Oct4, and Sox2 (Ding et al. 2015), also induces apoptosis during *in vitro* PGC specification and likely *in vivo* germline development. And like Nanog (Murakami et al. 2016), ectopic expression of Tex10 greatly enhances PGC specification. Taken together, our data support the dual roles of the core ESC transcription factors in regulating both stem cell pluripotency and germ cell development and suggest that the disruption of the core pluripotency network would lead to PGC loss. However, due to the early embryonic lethality and the loss of stem cell maintenance often associated with the loss-of-functions of these factors (e.g., Nanog (Mitsui et al. 2003), Oct4 (Nichols et al. 1998), Sox2 (Avilion et al. 2003), and Tex10 (Ding et al. 2015)), understanding of their functional roles in germline development often requires an inducible or conditional KD/KO system. In this regard, our newly developed Tex10 degron ESCs and the conditional Tex10 KO mouse model enabled us to uncover the critical roles of Tex10 in PGC specification and spermatogenesis in this study and will provide valuable resources for future studies of Tex10 in other developmental contexts.

Tex10 was initially identified as an interacting partner of Sox2 in maintaining mouse ESC pluripotency (Ding et al. 2015). Conditional Sox2 KO in PGCs using *Prdm1*-Cre and *TNAP*-Cre deleter mice results in a dramatic decrease of germ cell numbers at the time of their specification and compromised PGC proliferation at a later stage, respectively (Campolo et al. 2013). It should be noted that both *Prdm1*-Cre and *TNAP-Cre-mediated Sox2-homozygous* KO results in early postnatal death of pups due to unintended deletion of Sox2 in other cells/tissues than the germline (Campolo et al. 2013). Despite its wide use in studying *in vivo* PGC development, *Prdm1*-Cre was reported to also be active in primitive endoderm (PrE) at E6.75-9.0 (Mikedis and Downs 2017), which overlapped with mouse PGC specification (E6.25-8) and migration (E8.5-10.5) stages *in vivo* (von Meyenn and Reik 2015). The yolk sac, which develops from the PrE, provides gas and nutrition exchange between the developing embryo and the mother (Donovan and Bordoni 2021). Taking into account the apoptosis induced by the depletion of Tex10 from our *in vitro* study (Fig. 2A, S2A and S2F) and the detectable expression of *Tex10* in extraembryonic tissues (Zhang et al. 2018b; Nowotschin et al. 2019), we speculated that the unintended knockout of Tex10 after *Prdm1*-Cre excision in extraembryonic tissues (yolk sac and PrE) could have compromised embryogenesis at E6.75-9.0. The reported unintended deletion of Sox2 in other cells/tissues than the germline (Campolo et al. 2013) also discouraged us from using *TNAP*-Cre in our current study.

*T* (also known as Brachyury) is a target gene of the Wnt/β-catenin signaling pathway (Arnold et al. 2000) and a key PGC gene (Aramaki et al. 2013). However, *T* is not a direct Tex10-binding target and is only slightly activated upon Tex10 depletion with aberrant activation of Wnt signaling (Fig. S2J). Furthermore, the potential positive roles of T activation in PGC specification, if any, can be negated by marked up-regulation of the PGC restriction factor Otx2 in cells depleted with Tex10 (Fig. S2J). It should also be noted that the depletion of Tex10 has affected only the Wnt signaling, but not the Bmp signaling (Fig. 2D). Therefore, Tex10 depletion likely compromises PGC development through a mechanism distinct from the classic Bmp4-induced signaling pathway, where activation of T and repression of Otx2 are typically observed (Yao et al. 2022). We speculate that the regulating effect of Tex10 on Wnt signaling in PGC specification is more likely due to its impact on direct Tex10 targets, notably the two Wnt negative regulators *Psmd3* and *Psmd7*, rather than on *T*, although how Tex10 may directly or indirectly restrict Otx2 and somatic lineage programs requires future investigation (Fig. S7H).

Finally, although we have identified direct transcriptional targets of Tex10 (i.e., *Psmd3/7*) for the negative regulation of Wnt signaling, it remains to be determined how extensive genome reprogramming and alteration of DNA methylation and histone modification (Nikolic et al. 2016) may contribute to transcriptional functions of Tex10. The enrichment of GO term “DNA methylation” in EpiLC and D2-4 PGCLC stages upon Tex10 depletion (Fig. S5) is particularly intriguing, considering that PGC specification is anti-correlated with DNA methylation (Kurimoto et al. 2008) and that Tex10 can bridge the core pluripotency factor Sox2 with DNA hydroxylase Tet1 for the DNA hypomethylation status of the super-enhancers associated with pluripotency genes (Ding et al. 2015). Future studies to map the Tex10 interactome in PGCs will reveal its potential physical and functional interactions with other PGC specifiers such as Oct4 and Nanog and epigenetic regulators such as Tet family proteins, unraveling the transcriptional and epigenetic roles of Tex10 in PGC specification and male germline development.

## Acknowledgments

We thank a shared instrumentation grant for the Influx (S10OD020056) and LSRII (S10RR027050) flow cytometer at the Columbia Center for Translational Immunology (CCTI) flow cytometry core. The research in the Wang laboratory is supported by NIH (GM129157, HD095938, and HD097268) and NYSTEM (C32583GG and C32569GG). This research used the Shared Resource for Genomics and High Throughput Screening funded partly through the NIH/NCI Cancer Center Support Grant P30CA013696 and the NIH National Center for Advancing Translational Sciences Grant UL1TR001873. The content is solely the responsibility of the authors and does not necessarily represent the official views of the NIH.

## Authors’ Contributions

J.Y. and F.M. designed the experiments. J.Y., F.M., D.L., R.Z., and X.S. performed the experiments. J.Y., V.M., and X.H. performed the computational analysis. J.Y. wrote the manuscript draft. H.Z. provided reagents and experimental support. J.W. conceived, designed, supervised the studies, and wrote and approved the final manuscript.

## MATERIALS AND METHODS

### Animals

As the Tex10 conventional knockout mice model is early embryonic lethal (Ding et al. 2015), we made conditional knockout mice for Tex10 to evaluate its function in germ cell development. Tex10 is a gene with multiple exons and has a few splicing isoforms in the first three exons. To preserve the internal initiation site and minimize the chance of downstream in-frame methionine codons to be used as translational start and produce a truncated protein, we only considered floxing the exon 5 and/or downstream exons. Given the convenience of genotyping with only 113 bp (the coding sequence of exon 5), the exon 5 was chosen to be excised to functionally disrupt the Tex10 gene. Accordingly, two loxP sites were inserted to flank the exon 5. Such deletion would produce transcripts with premature stop codons and activated nonsense-mediated mRNA decay, thus leading to Tex10 knockout. However, as loxP is only 34 bp in length, it is not easy to distinguish the wildtype allele and the inserted allele. To overcome this hurdle, a BamHI site was also added to loxP. Then, Cre-Lox recombination was used to delete the DNA fragment between these two loxP sites (Fig. S6A).

CRISPR/Cas9 targeting reagents for the Tex10 conditional knockout (KO) founder lines were designed, generated and confirmed or validated at the Genome Engineering & iPSC Center (GEiC) at Washington University (WashU St. Louis, MO). These reagents were delivered to the Mouse Genetics and Gene Targeting (MGGT) core in the Icahn School of Medicine at Mount Sinai for zygote injection to generate CRISPR babies, among which the founder mice were identified. By co-injecting Cas9 plasmid, validated gRNAs, and single-strand oligonucleotide donor DNAs (ssODN) into fertilized eggs from C57BL/6N mice, we obtained 66 mice (one died shortly after being born) for next-gen sequencing (NGS service provided by WashU St. Louis, MO) and identified three mice with correct loxP-BamHI sites in both introns 4 and 5 of the same allele, namely founders #45 (male), #46 (male), #53 (female). To ensure that loxP-BamHI sites can be transmitted to the next generation, the three founders were mated with wild-type C57BL/6N mice and the resulting pups were genotyped for 5’ and 3’ loxP-BamHI sites, respectively. Fragments containing the two loxP-BamHI sites were amplified by F1∼F4 (refer to Table 16) and sequenced again for verification. The pups from founder #45 (male) had the correct loxP-BamHI sites and intact introns 4 and 5, which are used for subsequent studies. Three primers, namely F1, F2, and F4 (Fig. S6A and the Table S16), were designed for genotyping. The deletion by certain Cre strains produced the “-” band at 360 bp from the 470-bp “f” loxP band, and the wildtype “+” was at 430 bp (Fig. S6A). The gRNA sequences for intron 4: 5’-cctatgggctgggccctacataa-3’ and for intron 5: 5’-ccataagtgtggcacctgtgtgc-3’.

All animal experiments were conducted according to institutional guidelines for animal welfare and approved by the Institutional Animal Care and Use Committee (IACUC) of Columbia University. Embryos were isolated from 8-week-old female mice on the indicated day after fertilization. The testes were isolated from 8-week-old male mice. The ovaries were isolated from 28-day female mice.

### Cell lines

Male J1 mouse ESCs (mESCs) were maintained in an ES medium containing 80% high-glucose DMEM, 15% fetal bovine serum (FBS), 1% nucleoside mix, 100 μM nonessential amino acids, 50 U/mL penicillin/streptomycin, 0.1 mM 2-mercaptoethanol, 2 mM L-glutamine, and homemade recombinant leukemia inhibitory factor (LIF).

### Induction of EpiLCs and PGCLCs

EpiLCs and PGCLCs were induced as previously described (Hayashi et al. 2011). Briefly, mESCs were cultured in 2i (3 μM CHIR99021 and 1 μM PD0325901) plus LIF medium for seven days to achieve the naïve pluripotency state. The medium was then replaced with FA (12 ng/mL bFGF and 20 ng/mL Activin A) medium for two days to obtain EpiLCs. After that, 5 × 10^4^ EpiLCs were plated in one well of 24-well plates with Pluronic® F-127 coating to make it low attachment, thus preventing cell adhesion on the bottom of wells (Treter et al. 2014). PGCLC medium containing the cytokines BMP4 (500 ng/mL), LIF (1000 u/ml), SCF (100 ng/mL), BMP8b (500 ng/mL), and EGF (50 ng/mL) was used for six days, and cells were used to detect specification efficiency by flow cytometry. For cells with Tex10 overexpression, PGCLC induction was performed in the presence and absence of these cytokines for comparison.

### FACS

PGCLCs were dissociated with 0.05% Trypsin-EDTA (GIBCO), washed with DMEM containing 10% FBS, and resuspended with 1×PBS containing 0.1% BSA. Large cell clumps were removed using a cell strainer (Falcon™ 352235). The cells were analyzed and sorted on flow cytometers.

### Staining for FACS

Following induction of EpiLCs and PGCLCs as previously described, after two days of incubation with PGCLC medium, cells were stained with SSEA1 and CD61 for 30 min and then washed with MACS buffer (1% BSA, 1 mM EDTA in PBS). Cells were collected by centrifugation at 400 g for 5 min at 4 °C and fixed with 4% PFA for 10 min. The cells were then washed with PBS and permeabilized with cold 0.1% Triton-X-100 in PBS (PBST) on ice for 10 min. After washing with PBST, cells were blocked with 2% BSA in PBST at room temperature for 30 min. Cells were then washed with PBST and resuspended in MACS buffer with the primary antibody (Prdm1, Cell Signaling, 9115S) for 1.5 hours at 4 °C in the dark. The cells were then washed with PBST and stained with secondary antibody for 30 min at 4 °C. Next, after washing with PBST and then MACS buffer, the cells were resuspended with MACS buffer, and large clumps of cells were removed using a cell strainer (Falcon™ 352235). Finally, cells were analyzed on flow cytometers.

### Preparation of adult murine testicular cell suspension, Hoechst dye staining, flow cytometric analysis, and single-cell sequencing

Mouse testicular cell suspension, Hoechst dye staining, and flow cytometric analysis were performed following the published protocol (Gaysinskaya et al. 2014). Briefly, testicular tissues were digested with 7 ml collagenase type IV (290U/ml, Gibco cat # 17104-019) for 10 min at 37 °C with gentle agitation (250 rpm) in 15 ml centrifuge tubes. The tubules were naturally sedimented for 1 min, and 6 ml suspension was discarded. Four ml of PBS, 5 ml 0.25% trypsin/EDTA (Gibco cat:25200-056), and 10 μg/ml of DNase I were added for digestion for 8 min at 37 °C. Six hundred μL FBS was used to end digestion. For single cell analysis, the above cells were filtered with 70 μm (Falcon cat #352350) and 40 μm (Falcon cat #352340) cell strainer, and the filtered cells were used for single-cell sequencing in the Columbia University genomic core facility. Briefly, 10x Genomics Chromium Single Cell 3’ Reagent Kits v3.1 Chemistry with Dual Indexing were used for library preparation according to the manufacturer’s instructions, and the NovaSeq 6000 was used for sequencing (2×100bp). For flow cytometric analysis, cells were also filtered with 70 μm (Falcon cat #352350) cell strainer and treated with 100 μg Hoechst for 20 min at room temperature (RT). These pre-stained cells were centrifuged and resuspended with DMEM supplemented with 10% FBS and 10 μg/ml DNase I. After this, cells were stained with 6 μg Hoechst per million cells for 30 min at RT. Stained cells were filtered with a cell strainer (Falcon Cat#352235) and put on ice until flow cytometric analysis. Samples were analyzed by LSRII flow cytometry in the Columbia University flow core facility.

### RT-qPCR

Total RNAs from EpiLCs and PGCLCs were extracted using Trizol (Fisher, 15596018) and converted to cDNA using qScript (Quanta). Relative gene expression levels were analyzed with PowerUp™ SYBR™ Green Master Mix on the QuantStudio 5 PCR system (Life Technologies Inc.). Gene expression levels were normalized to *Gapdh*.

### Western blot

Cells were lysed in RIPA buffer (Boston BioProducts, Inc.) supplied with PMSF (Roche, 11359061001) and protease inhibitor cocktail (Sigma, P8340). Protein concentration was measured by Bradford assay reagent (Thermo Scientific). The protein amounts were adjusted among samples, then 4 × LDS sample buffer (GenScript, M00676-10) was added. Samples were boiled at 95 °C for 5 min. Proteins were separated on GenScript SurePAGETM Bis-Tris gels and blotted on the Immobilon-P transfer membrane (Millipore). The membrane was blocked with 5% skim milk and incubated with primary antibodies: anti-Nanog (1:500, rabbit IgG, Bethyl/Fisher, A300-397A), anti-Gapdh (1:1000, rabbit IgG, ProteinTech, 10494-1-AP), anti-HA tag (1:1000, rabbit IgG, Abcam, ab9110), anti-β-actin (1:1000, mouse IgG, Sigma-Aldrich, clone AC-15, A5441). anti-Otx2 (1:1000, rabbit IgG, Abcam, ab21990), anti-Prdm1 (1:1,000, mouse IgG, Sigma-Aldrich, clone 5E7, SAB5300402), anti-Tex10 (1:1000, rabbit IgG, Thermo Fisher, 720257), anti-Vinculin (1:10000, rabbit IgG, Abcam, ab129002), anti-Psmd7 (1:1000, mouse IgG, Santa Cruz, sc-390705). Then it was incubated with a secondary antibody and detected using a Medical Film Processor (SRX-101A) or ImageQuant LAS 4000 (GE Healthcare).

### Immunofluorescence staining of embryoid bodies

Embryoid bodies (EBs) were fixed with 4% paraformaldehyde (w/v) for 15 min at RT, then washed and permeabilized with 0.25% Triton X-100 solution for 5 min at RT. After that, they were blocked with 5% fetal bovine serum (FBS) in PBS. Primary antibodies were incubated with EBs overnight at 4°C. The next day, the EBs were washed and incubated with secondary antibodies and 3 μM DAPI for 1 hour at RT in the dark. After washing, the EBs were imaged with a Leica DMi8 inverted microscope.

### Transfection and lentiviral infection

Transfection of cells was performed using lipofectamine 3000 according to the manufacturer’s instructions. The production of lentivirus and viral infection were performed as described (Ivanova et al. 2006). The Psmd7 plasmid used for ectopic overexpression was purchased from NovoPro (Cat. 749214-1)

### Histology

The ovaries were first fixed in 4% paraformaldehyde at 4°C overnight. Then, they were dehydrated through a series of grade ethanol and embedded in paraffin. Finally, sections were performed and mounted on the slides for Hematoxylin and Eosin (H&E) staining.

### Immunofluorescence staining of paraffin sections

The paraffin sections were dewaxed and rehydrated. Antigen retrieval was performed by high-pressure heating with a commercial antigen unmasking retrieval solution and then blocked with 5% fetal bovine serum. Sections were incubated with primary antibodies anti-β-galactosidase (1:500, mouse IgG, Thermo Fisher, Z3781), anti-Sox9 (1:1000, rabbit IgG, Sigma-Aldrich, AB5535) at 4°C overnight. Conjugated secondary antibodies (1:500) were added to the sections and incubated for 2 h at RT. The nucleus was stained with Hoechst 33342, and images were obtained with a Leica DMI 6000 inverted microscope.

### LacZ staining

The ovary was fixed, washed, and then incubated with X-gal staining solution overnight at 37°C. After two washes in PBS, the sample was post-fixed with 4% paraformaldehyde at 4°C for 24 h. Following two washes in PBS, the sample was used for paraffin sectioning and counterstaining.

### Degron system plasmid construction

The 5’ arm and 3’arm of *Tex10* C-terminal were cloned from the genomic DNA of J1 mESCs, and FKBP-V-P2A-BFP was cloned from pAW63.YY1.FKBP.knock-in.BFP plasmid (Addgene plasmid #104371). Then, Gibson cloning was conducted to connect the above three fragments. The PCR product from the Gibson assembly was purified and cloned into pJet1.2 plasmid (CloneJET PCR Cloning Kit, ThermoFisher). Then the loxP-dsRed-loxP fragment was cloned from pMSCV-loxP-dsRed-loxP-3xHA-Puro-WPRE (Addgene plasmid #32703). The loxP-dsRed-loxP fragment and the above modified pJet1.2 plasmid were digested with the restriction enzymes AgeI (NEB, R0552S) and PacI (NEB, R0547S) and then ligated using T4 DNA ligase (NEB, M0202S) to obtain the plasmid pAW63-Tex10. The Tex10 guide RNA sequence (refer to Table S16) was inserted into the PX330-puro plasmid (Addgene plasmid #62988) to obtain the PX330-Tex10 plasmid.

### RNA-seq

J1 mESCs were transfected with plasmids pAW63-Tex10 and PX330-Tex10 using Lipofectamine 3000 (Thermo Fisher, L3000001) according to the manufacturer’s manual, followed by drug selection with puromycin (1μg/mL) for three days. The selected cells were transfected with the pPyCAG-CreERT2-ires-Puro plasmid, modified from pPyCAG-Cre::ERT2-IRES-BSD (Addgene plasmid #48760) by changing blasticidin-resistance gene to puromycin-resistance gene, and then treated with 4-hydroxytamoxifen (Sigma, H7904) to delete the dsRed fragment. Clones without fluorescence were picked and expanded. Expanded clones were treated with or without dTAG13 (500 nM, Toronto Research Chemicals, D710020) and compared with wildtype J1 mESCs for Tex10 expression. The size difference of Tex10 protein between successful clones and wildtype is 12 kDa, which is the size of FKBP12^F36V^, and dTAG13 treatment (500 nM) fully degraded the Tex10 protein. Two clones were validated and used for the following *in vitro* PGCLC specification experiments. Biological duplicates were prepared for cells at EpiLC, D2, D4, and D6 PGCLC stages. EBs treated with dTAG13 for two (dD2EBs) and four days (dD4EBs) were collected, and EBs treated with DMSO for two (D2EBs) and four days (D4EBs) were also collected. Too few cells survived after dTAG13 treatment for six days, so they were not collected. Total RNA was extracted with Trizol, and then RNA-seq libraries were made with 100 ng RNA using the Ovation Mouse RNA-seq kit (NuGEN, #0348-32) according to the manufacturer’s protocol. Tex10 overexpression and control mESCs were transfected with the FLAG-Tex10 plasmid and the empty vector (EV) plasmid, respectively (Ding et al. 2015). Biological duplicates were prepared for Tex10 overexpression cells. The overexpression was first confirmed by western blotting analysis. Then, confirmed cells were used for *in vitro* PGC specification. Cells at EpiLC, D2, D4, and D6PGCLC stages were collected. Total RNA was extracted using Trizol and followed by library preparation using Universal Plus mRNA-Seq with the NuQuant Kit (TECAN, 0520-A01). Mouse testis samples were extracted from testes of Tex10 cKO (*Tex10* f/f;*Stra8*-iCre) and WT (*Tex10* f/f) mice, followed by RNA extraction using Trizol and library preparation using the Ovation Mouse RNA-seq kit (NuGEN, #0348-32). Libraries were sequenced on the HiSeq 4000 to obtain pairedend 150-nucleotide read length.

### Chromatin immunoprecipitation (ChIP)

ChIP-qPCR and ChIP-seq samples were prepared following the instructions of the SampleChIP^R^ Enzymatic Chromatin IP Kit (Cell Signaling, #9005). Briefly, 4 × 10^6^ cells per ChIP were fixed in 1% formaldehyde (RT, 10 min), quenched with 1 volume of 250 mM glycine (RT, 5 min), and rinsed twice with chilled PBS buffer before storage at −80 °C. After thawing cells on ice, fixed cells were lysed to pellet nuclei and 0.5 μl micrococcal nuclease per sample was used to digest DNA. The digestion was stopped by adding 10 μl 0.5 M EDTA per sample and placed on ice for 2 min. The nuclei pellet was then collected by 16,000 g at 4 °C for 1 min. The pellets of the individual samples were resuspended in 100 μl 1X ChIP buffer and incubated on ice for 10 min. The samples were sonicated five times (30-s pulses with 30-s break interval) using the Bioruptor water bath sonicator (Diagenode). The chromatin extracts were then centrifuged at 9,400 g for 10 min at 4 °C. The supernatant was transferred to a new tube for the following IP experiments. For each sample, 100 μl digested chromatin was diluted into 400 μl 1X ChIP buffer. We used two cell clones, C1 and C2, as biological replicates, and 10 μl of diluted chromatin was used as 2% input. For each IP, anti-HA tag antibody (6 μg per ChIP, Abcam, ab9110), H3K4me3 antibody (6 μg per ChIP, Abcam, ab8580), or normal rabbit IgG (6 μg per ChIP, Cell Signaling Tech., 2729P) as a negative control was incubated with digested chromatin overnight with rotation at 4 °C and then incubated with 30 μl Protein G magnetic beads for 2 h at 4 °C with rotation. Using the magnetic separation rack, the beads were washed with low-salt buffer three times and high-salt buffer one time at 4 °C with rotation. For input samples, 150 μl ChIP elution buffer was added and put aside at RT. For IP samples, 150 μl ChIP elution buffer was added and incubated at 65 °C for 30 min with gentle vortexing. A magnetic separation rack was used to get the solution. Reverse crosslinking was performed at 65 °C for 6 h by adding 2 μl Proteinase K and 6 μl 5M NaCl. Finally, DNA was purified for qPCR analysis or used for library preparation using NEBNext® Ultra™ II DNA Library Prep (E7103S). Once prepared, the library was sequenced using HiSeq 4000 with paired-end 150 nucleotides read length.

### RNA-seq data analysis

RNA-seq reads were aligned to the mouse mm10 genome using Bowtie2 (v2.3.4.3), and aligned bam files were sorted by name using the parameter -n. Next, we used the HTSeq software (v0.11.2) and the GENCODE mm10 annotation file to count reads for each gene using the parameters -r name -f bam and BioMart (Durinck et al. 2005) to retrieve the names of the corresponding genes. Finally, read counts were normalized with the trimmed mean of M-values (TMM) method (Robinson and Oshlack 2010) for differential expression analysis using edgeR (v3.26.8) (Robinson et al. 2010). The public RNA-seq data were then downloaded (refer to the Table S16) and then aligned to mm10 using the same processing setting.

### Single-cell RNA-seq data analysis

Single cell RNA sequencing reads were aligned with the mm10 genome to generate an expression matrix using Cell Ranger (V4.0.0), and the produced data were further filtered using Seurat (V4.0.5) to exclude cells with fewer than 200 genes expressed and >50% of the total expression of mitochondrial genes. Through the above steps, 29224 cells were used for the following analysis, consisting of 17816 cells from WT and 11408 cells from the cKO mouse testis. To remove the batch effect, the top 2000 highly variable genes were used for the canonical correlation analysis implemented in Seurat (V4.0.5).

### ChIP-seq data analysis

ChIP-seq reads were mapped to the mm10 genome using Bowtie2 (v2.3.4.3), and PCR duplicates were removed using Samtools (v1.9). Then the peaks were called using MACS2 (v2.1.2) to compare the IP samples with the input samples, only the peaks called from both replicates were kept. Finally, annotation was done using the script annotatePeaks.pl of HOMER (Heinz et al. 2010).

### The gene set activity analysis

Pluripotency, PGC regulator, mesoderm, endoderm, and ectoderm gene sets were curated in the literature (Ding et al. 2015; Hackett et al. 2018) and in the R&D database (Table S15). First, for each gene in a specific category, such as pluripotency, the log_2_FC of the gene was calculated. Then the mean log_2_FC of all genes in a category was considered as the overall activity value.

## QUANTIFICATION AND STATISTICAL ANALYSIS

Statistical analyses were carried out using R (version 3.5.2) or Prism (version 9.1.2). Data are presented as mean ± SD. The student’s *t-*test was used to determine the statistical significance if not stated in the relevant figure legends. The criteria were applied to Figs. S1C, 6C, 6D, S6K, S7F, and S7G. **** for p-value < 0.0001, *** for p-value < 0.001, ** for p-value < 0.01, * for p-value < 0.05, and ns for p-value > 0.05.

## DATA AND CODE AVAILABILITY

All high-throughput ChIP-seq and RNA-seq data generated in this study are available on Gene Expression Omnibus under accession codes GSE180801 and GSE190072. To review GEO accession GSE180801: Go to https://www.ncbi.nlm.nih.gov/geo/query/acc.cgi?acc=GSE180801, enter token “epsjssmqnvuflep” into the box; to review GEO accession GSE190072: Go to https://www.ncbi.nlm.nih.gov/geo/query/acc.cgi?acc=GSE190072, enter token “clsbaywydbaxlij” into the box.

## REFERENCES

Aberle H, Bauer A, Stappert J, Kispert A, Kemler R. 1997. beta-catenin is a target for the ubiquitin-proteasome pathway. EMBO J 16: 3797–3804.

Aramaki S, Hayashi K, Kurimoto K, Ohta H, Yabuta Y, Iwanari H, Mochizuki Y, Hamakubo T, Kato Y, Shirahige K et al. 2013. A mesodermal factor, T, specifies mouse germ cell fate by directly activating germline determinants. Dev Cell 27: 516–529.

Arnold SJ, Stappert J, Bauer A, Kispert A, Herrmann BG, Kemler R. 2000. Brachyury is a target gene of the Wnt/beta-catenin signaling pathway. Mech Dev 91: 249–258.

Avilion AA, Nicolis SK, Pevny LH, Perez L, Vivian N, Lovell-Badge R. 2003. Multipotent cell lineages in early mouse development depend on SOX2 function. Genes Dev 17: 126–140.

Beke A. 2019. Genetic Causes of Female Infertility. Exp Suppl 111: 367–383.

Boyer LA, Lee TI, Cole MF, Johnstone SE, Levine SS, Zucker JP, Guenther MG, Kumar RM, Murray HL, Jenner RG et al. 2005. Core transcriptional regulatory circuitry in human embryonic stem cells. Cell 122: 947–956.

Broughton DE, Moley KH. 2017. Obesity and female infertility: potential mediators of obesity’s impact. Fertil Steril 107: 840–847.

Campolo F, Gori M, Favaro R, Nicolis S, Pellegrini M, Botti F, Rossi P, Jannini EA, Dolci S. 2013. Essential role of Sox2 for the establishment and maintenance of the germ cell line. Stem Cells 31: 1408–1421.

Chassot AA, Le Rolle M, Jourden M, Taketo MM, Ghyselinck NB, Chaboissier MC. 2017. Constitutive WNT/CTNNB1 activation triggers spermatogonial stem cell proliferation and germ cell depletion. Dev Biol 426: 17–27.

Chen D, Gell JJ, Tao Y, Sosa E, Clark AT. 2017. Modeling human infertility with pluripotent stem cells. Stem Cell Res 21: 187–192.

Chen SR, Batool A, Wang YQ, Hao XX, Chang CS, Cheng CY, Liu YX. 2016. The control of male fertility by spermatid-specific factors: searching for contraceptive targets from spermatozoon’s head to tail. Cell death & disease 7: e2472.

Craig JR, Jenkins TG, Carrell DT, Hotaling JM. 2017. Obesity, male infertility, and the sperm epigenome. Fertil Steril 107: 848–859.

de Vries WN, Binns LT, Fancher KS, Dean J, Moore R, Kemler R, Knowles BB. 2000. Expression of Cre recombinase in mouse oocytes: a means to study maternal effect genes. Genesis 26: 110–112.

Ding J, Huang X, Shao N, Zhou H, Lee DF, Faiola F, Fidalgo M, Guallar D, Saunders A, Shliaha PV et al. 2015. Tex10 Coordinates Epigenetic Control of Super-Enhancer Activity in Pluripotency and Reprogramming. Cell Stem Cell 16: 653–668.

Donovan MF, Bordoni B. 2021. Embryology, Yolk Sac. in StatPearls, Treasure Island (FL).

Durinck S, Moreau Y, Kasprzyk A, Davis S, De Moor B, Brazma A, Huber W. 2005. BioMart and Bioconductor: a powerful link between biological databases and microarray data analysis. Bioinformatics 21: 3439–3440.

Fang X, Huang LL, Xu J, Ma CQ, Chen ZH, Zhang Z, Liao CH, Zheng SX, Huang P, Xu WM et al. 2019. Proteomics and single-cell RNA analysis of Akap4-knockout mice model confirm indispensable role of Akap4 in spermatogenesis. Dev Biol 454: 118–127.

Fararjeh AS, Chen LC, Ho YS, Cheng TC, Liu YR, Chang HL, Chang HW, Wu CH, Tu SH. 2019. Proteasome 26S Subunit, non-ATPase 3 (PSMD3) Regulates Breast Cancer by Stabilizing HER2 from Degradation. Cancers (Basel) 11.

Fawcett DW, Phillips DM. 1969. The fine structure and development of the neck region of the mammalian spermatozoon. Anat Rec 165: 153–164.

Festuccia N, Osorno R, Wilson V, Chambers I. 2013. The role of pluripotency gene regulatory network components in mediating transitions between pluripotent cell states. Curr Opin Genet Dev 23: 504–511.

Gaysinskaya V, Soh IY, van der Heijden GW, Bortvin A. 2014. Optimized flow cytometry isolation of murine spermatocytes. Cytometry Part A: the journal of the International Society for Analytical Cytology 85: 556–565.

Hackett JA, Huang Y, Gunesdogan U, Gretarsson KA, Kobayashi T, Surani MA. 2018. Tracing the transitions from pluripotency to germ cell fate with CRISPR screening. Nature communications 9: 4292.

Hayashi K, Ohta H, Kurimoto K, Aramaki S, Saitou M. 2011. Reconstitution of the mouse germ cell specification pathway in culture by pluripotent stem cells. Cell 146: 519–532.

Heinz S, Benner C, Spann N, Bertolino E, Lin YC, Laslo P, Cheng JX, Murre C, Singh H, Glass CK. 2010. Simple combinations of lineage-determining transcription factors prime cis-regulatory elements required for macrophage and B cell identities. Molecular cell 38: 576–589.

Hemendinger RA, Gores P, Blacksten L, Harley V, Halberstadt C. 2002. Identification of a specific Sertoli cell marker, Sox9, for use in transplantation. Cell Transplant 11: 499–505.

Hermann BP, Cheng K, Singh A, Roa-De La Cruz L, Mutoji KN, Chen IC, Gildersleeve H, Lehle JD, Mayo M, Westernstroer B et al. 2018. The Mammalian Spermatogenesis Single-Cell Transcriptome, from Spermatogonial Stem Cells to Spermatids. Cell reports 25: 1650–1667 e1658.

Hutchins AP, Yang Z, Li Y, He F, Fu X, Wang X, Li D, Liu K, He J, Wang Y et al. 2017. Models of global gene expression define major domains of cell type and tissue identity. Nucleic Acids Res 45: 2354–2367.

Ivanova N, Dobrin R, Lu R, Kotenko I, Levorse J, DeCoste C, Schafer X, Lun Y, Lemischka IR. 2006. Dissecting self-renewal in stem cells with RNA interference. Nature 442: 533–538.

Kashiwabara S, Noguchi J, Zhuang T, Ohmura K, Honda A, Sugiura S, Miyamoto K, Takahashi S, Inoue K, Ogura A et al. 2002. Regulation of spermatogenesis by testis-specific, cytoplasmic poly(A) polymerase TPAP. Science 298: 1999–2002.

Kehler J, Tolkunova E, Koschorz B, Pesce M, Gentile L, Boiani M, Lomeli H, Nagy A, McLaughlin KJ, Scholer HR et al. 2004. Oct4 is required for primordial germ cell survival. EMBO Rep 5: 1078–1083.

Kimura T, Kaga Y, Ohta H, Odamoto M, Sekita Y, Li K, Yamano N, Fujikawa K, Isotani A, Sasaki N et al. 2014. Induction of primordial germ cell-like cells from mouse embryonic stem cells by ERK signal inhibition. Stem Cells 32: 2668–2678.

Kominami K, Okura N, Kawamura M, DeMartino GN, Slaughter CA, Shimbara N, Chung CH, Fujimuro M, Yokosawa H, Shimizu Y et al. 1997. Yeast counterparts of subunits S5a and p58 (S3) of the human 26S proteasome are encoded by two multicopy suppressors of nin1-1. Mol Biol Cell 8: 171–187.

Krausz C, Riera-Escamilla A. 2018. Genetics of male infertility. Nat Rev Urol 15: 369–384.

Kurimoto K, Yabuta Y, Ohinata Y, Shigeta M, Yamanaka K, Saitou M. 2008. Complex genome-wide transcription dynamics orchestrated by Blimp1 for the specification of the germ cell lineage in mice. Genes Dev 22: 1617–1635.

Liberzon A, Birger C, Thorvaldsdottir H, Ghandi M, Mesirov JP, Tamayo P. 2015. The Molecular Signatures Database (MSigDB) hallmark gene set collection. Cell Syst 1: 417–425.

Liu D, Liao C, Wolgemuth DJ. 2000. A role for cyclin A1 in the activation of MPF and G2-M transition during meiosis of male germ cells in mice. Dev Biol 224: 388–400.

Liu D, Matzuk MM, Sung WK, Guo Q, Wang P, Wolgemuth DJ. 1998. Cyclin A1 is required for meiosis in the male mouse. Nat Genet 20: 377–380.

Liu S, Brind’Amour J, Karimi MM, Shirane K, Bogutz A, Lefebvre L, Sasaki H, Shinkai Y, Lorincz MC. 2014. Setdb1 is required for germline development and silencing of H3K9me3-marked endogenous retroviruses in primordial germ cells. Genes Dev 28: 2041–2055.

Magnusdottir E, Dietmann S, Murakami K, Gunesdogan U, Tang F, Bao S, Diamanti E, Lao K, Gottgens B, Azim Surani M. 2013. A tripartite transcription factor network regulates primordial germ cell specification in mice. Nat Cell Biol 15: 905–915.

Mikedis MM, Downs KM. 2017. PRDM1/BLIMP1 is widely distributed to the nascent fetal-placental interface in the mouse gastrula. Dev Dyn 246: 50–71.

Mitsui K, Tokuzawa Y, Itoh H, Segawa K, Murakami M, Takahashi K, Maruyama M, Maeda M, Yamanaka S. 2003. The homeoprotein Nanog is required for maintenance of pluripotency in mouse epiblast and ES cells. Cell 113: 631–642.

Mitsunaga K, Araki K, Mizusaki H, Morohashi K, Haruna K, Nakagata N, Giguere V, Yamamura K, Abe K. 2004. Loss of PGC-specific expression of the orphan nuclear receptor ERR-beta results in reduction of germ cell number in mouse embryos. Mech Dev 121: 237–246.

Mochizuki K, Hayashi Y, Sekinaka T, Otsuka K, Ito-Matsuoka Y, Kobayashi H, Oki S, Takehara A, Kono T, Osumi N et al. 2018. Repression of Somatic Genes by Selective Recruitment of HDAC3 by BLIMP1 Is Essential for Mouse Primordial Germ Cell Fate Determination. Cell reports 24: 2682–2693 e2686.

Murakami K, Gunesdogan U, Zylicz JJ, Tang WWC, Sengupta R, Kobayashi T, Kim S, Butler R, Dietmann S, Surani MA. 2016. NANOG alone induces germ cells in primed epiblast in vitro by activation of enhancers. Nature 529: 403–407.

Nabet B, Roberts JM, Buckley DL, Paulk J, Dastjerdi S, Yang A, Leggett AL, Erb MA, Lawlor MA, Souza A et al. 2018. The dTAG system for immediate and target-specific protein degradation. Nat Chem Biol 14: 431–441.

Nakaki F, Hayashi K, Ohta H, Kurimoto K, Yabuta Y, Saitou M. 2013. Induction of mouse germ-cell fate by transcription factors in vitro. Nature 501: 222–226.

Nichols J, Zevnik B, Anastassiadis K, Niwa H, Klewe-Nebenius D, Chambers I, Scholer H, Smith A. 1998. Formation of pluripotent stem cells in the mammalian embryo depends on the POU transcription factor Oct4. Cell 95: 379–391.

Nikolic A, Volarevic V, Armstrong L, Lako M, Stojkovic M. 2016. Primordial Germ Cells: Current Knowledge and Perspectives. Stem cells international 2016: 1741072.

Nowotschin S, Setty M, Kuo YY, Liu V, Garg V, Sharma R, Simon CS, Saiz N, Gardner R, Boutet SC et al. 2019. The emergent landscape of the mouse gut endoderm at single-cell resolution. Nature 569: 361–367.

Ohinata Y, Payer B, O’Carroll D, Ancelin K, Ono Y, Sano M, Barton SC, Obukhanych T, Nussenzweig M, Tarakhovsky A et al. 2005. Blimp1 is a critical determinant of the germ cell lineage in mice. Nature 436: 207–213.

Poorvu PD, Frazier AL, Feraco AM, Manley PE, Ginsburg ES, Laufer MR, LaCasce AS, Diller LR, Partridge AH. 2019. Cancer Treatment-Related Infertility: A Critical Review of the Evidence. JNCI Cancer Spectr 3: pkz008.

Rattner A, Hsieh JC, Smallwood PM, Gilbert DJ, Copeland NG, Jenkins NA, Nathans J. 1997. A family of secreted proteins contains homology to the cysteine-rich ligand-binding domain of frizzled receptors. Proc Natl Acad Sci U S A 94: 2859–2863.

Robinson MD, McCarthy DJ, Smyth GK. 2010. edgeR: a Bioconductor package for differential expression analysis of digital gene expression data. Bioinformatics 26: 139–140.

Robinson MD, Oshlack A. 2010. A scaling normalization method for differential expression analysis of RNA-seq data. Genome Biol 11: R25.

Sadate-Ngatchou PI, Payne CJ, Dearth AT, Braun RE. 2008. Cre recombinase activity specific to postnatal, premeiotic male germ cells in transgenic mice. Genesis 46: 738–742.

Salazar G, Joshi A, Liu D, Wei H, Persson JL, Wolgemuth DJ. 2005. Induction of apoptosis involving multiple pathways is a primary response to cyclin A1-deficiency in male meiosis. Dev Dyn 234: 114–123.

Subramanian A, Tamayo P, Mootha VK, Mukherjee S, Ebert BL, Gillette MA, Paulovich A, Pomeroy SL, Golub TR, Lander ES et al. 2005. Gene set enrichment analysis: a knowledge-based approach for interpreting genome-wide expression profiles. Proc Natl Acad Sci U S A 102: 15545–15550.

Takase HM, Nusse R. 2016. Paracrine Wnt/beta-catenin signaling mediates proliferation of undifferentiated spermatogonia in the adult mouse testis. Proc Natl Acad Sci U S A 113: E1489–1497.

Tang WW, Kobayashi T, Irie N, Dietmann S, Surani MA. 2016. Specification and epigenetic programming of the human germ line. Nat Rev Genet 17: 585–600.

Tezuka N, Brown AM, Yanagawa S. 2007. GRB10 binds to LRP6, the Wnt co-receptor and inhibits canonical Wnt signaling pathway. Biochemical and biophysical research communications 356: 648–654.

Tokue M, Ikami K, Mizuno S, Takagi C, Miyagi A, Takada R, Noda C, Kitadate Y, Hara K, Mizuguchi H et al. 2017. SHISA6 Confers Resistance to Differentiation-Promoting Wnt/beta-Catenin Signaling in Mouse Spermatogenic Stem Cells. Stem Cell Reports 8: 561–575.

Tokuhiro K, Isotani A, Yokota S, Yano Y, Oshio S, Hirose M, Wada M, Fujita K, Ogawa Y, Okabe M et al. 2009. OAZ-t/OAZ3 is essential for rigid connection of sperm tails to heads in mouse. PLoS genetics 5: e1000712.

Treter J, Bonatto F, Krug C, Soares GV, Baumvol IJR, Macedo AJ. 2014. Washing-resistant surfactant coated surface is able to inhibit pathogenic bacteria adhesion. Applied surface science 303: 147–154.

von Meyenn F, Reik W. 2015. Forget the Parents: Epigenetic Reprogramming in Human Germ Cells. Cell 161: 1248–1251.

Woolley DM. 2010. Flagellar oscillation: a commentary on proposed mechanisms. Biol Rev Camb Philos Soc 85: 453–470.

Yamaguchi S, Kurimoto K, Yabuta Y, Sasaki H, Nakatsuji N, Saitou M, Tada T. 2009. Conditional knockdown of Nanog induces apoptotic cell death in mouse migrating primordial germ cells. Development 136: 4011–4020.

Yang K, Meinhardt A, Zhang B, Grzmil P, Adham IM, Hoyer-Fender S. 2012. The small heat shock protein ODF1/HSPB10 is essential for tight linkage of sperm head to tail and male fertility in mice. Mol Cell Biol 32: 216–225.

Yao C, Yao R, Luo H, Shuai L. 2022. Germline specification from pluripotent stem cells. Stem Cell Res Ther 13: 74.

Yi JJ, Paranjape SR, Walker MP, Choudhury R, Wolter JM, Fragola G, Emanuele MJ, Major MB, Zylka MJ. 2017. The autism-linked UBE3A T485A mutant E3 ubiquitin ligase activates the Wnt/beta-catenin pathway by inhibiting the proteasome. J Biol Chem 292: 12503–12515.

Yuan S, Stratton CJ, Bao J, Zheng H, Bhetwal BP, Yanagimachi R, Yan W. 2015. Spata6 is required for normal assembly of the sperm connecting piece and tight head-tail conjunction. Proc Natl Acad Sci U S A 112: E430–439.

Zhang J, Zhang M, Acampora D, Vojtek M, Yuan D, Simeone A, Chambers I. 2018a. OTX2 restricts entry to the mouse germline. Nature 562: 595–599.

Zhang Y, Xiang Y, Yin Q, Du Z, Peng X, Wang Q, Fidalgo M, Xia W, Li Y, Zhao ZA et al. 2018b. Dynamic epigenomic landscapes during early lineage specification in mouse embryos. Nat Genet 50: 96–105.

Zheng M, Chen X, Cui Y, Li W, Dai H, Yue Q, Zhang H, Zheng Y, Guo X, Zhu H. 2021. TULP2, a New RNA-Binding Protein, Is Required for Mouse Spermatid Differentiation and Male Fertility. Frontiers in cell and developmental biology 9: 623738.

Zhuang T, Kashiwabara S, Noguchi J, Baba T. 2004. Transgenic expression of testis-specific poly(A) polymerase TPAP in wild-type and TPAP-deficient mice. J Reprod Dev 50: 207–213.

